# Genetic legacies of mega-landslides: cycles of isolation and contact across flank collapses in an oceanic island

**DOI:** 10.1101/2024.01.10.574994

**Authors:** Víctor Noguerales, Yurena Arjona, Víctor García-Olivares, Antonio Machado, Heriberto López, Jairo Patiño, Brent C. Emerson

## Abstract

Catastrophic flank collapses are recognised as important drivers of insular biodiversity dynamics, through the disruption of species ranges and subsequent allopatric divergence. However, little empirical data supports this conjecture, with their evolutionary consequences remaining poorly understood. Using genome-wide data within a population genomics and phylogenomics framework, we evaluate how mega-landslides have impacted evolutionary and demographic history within a species complex of weevils (Curculionidae) within the Canary island of Tenerife. We reveal a complex genomic landscape, within which individuals of single ancestry were sampled in areas characterised by long-term geological stability, relative to the timing of flank collapses. In contrast, individuals of admixed ancestry were almost exclusively sampled within the boundaries of flank collapses. Estimated divergence times among ancestral populations aligned with the timings of mega-landslide events. Our results provide first evidence for a cyclical dynamic of range fragmentation and secondary contact across flank collapse landscapes, with support for a model where this dynamic is mediated by Quaternary climate oscillations. The context within which we reveal climate and topography to interact cyclically through time to shape the geographic structure of genetic variation, together with related recent work, highlights the importance of topoclimatic phenomena as an agent of diversification within insular invertebrates.

## INTRODUCTION

Oceanic archipelagos are considered to serve as natural laboratories for evolutionary biologists and ecologists, providing an important framework to improve our understanding about the drivers of speciation (Losos & Rickkefs, 2009). Insular diversification is frequently associated with ecological gradients, but non-ecological mechanisms are also expected to promote speciation within insular settings by local geographic isolation (Goodman *et al*., 2012; Machado, 2022; Salces-Castellano *et al*., 2021). In this vein, it has been recently argued for the need to focus attention on geomorphological dynamics within islands, highlighting the role of volcanic eruptions and other major landform-changing events, such as mega-landslides, in the evolutionary process (Otto *et al*., 2016). During the typical developmental life cycle of oceanic islands, geological activity acts to remodel existing landscapes. This geological dynamic is represented by three major events: eruptive volcanic activity, catastrophic flank collapse and millennial-scale erosion. Of these three phenomena, the immediate consequences of eruptive events and flank collapses may directly impact island biotas by provoking local extinction in affected zones (Borges & Hortal, 2009). Such extinctions driven by volcanism (Bloor *et al*., 2008; Carson *et al*., 1990; Vandergast *et al*., 2004) or by landslides (Brown *et al*., 2006; Juan *et al*., 2000; Macías-Hernández *et al*., 2013) may be followed by long-term habitat discontinuities generating genetic differentiation among populations (Goodman *et al*., 2012). As conditions in the empty ecological space generated by a catastrophic geological event become suitable, recolonization from adjacent zones could promote episodes of secondary contact among previously isolated populations. Evidence for such a complex dynamic is limited, but several phylogeographic studies within the Canary Island of Tenerife provide some support (Figure S1). For instance, Brown *et al*. (2006) revealed the potential role of the Güímar flank collapse on cladogenesis through population fragmentation and isolation within the lizard *Gallotia galloti*, based on genetic and morphological data. Population differentiation within the spider species *Dysdera verneaui* coincides with the western and eastern division of the Anaga peninsula by the flanks of a mega-landslide (Macías-Hernández *et al*., 2013). An estimated landslide age of 0.5-1.0 million years (Ma) (Watts & Masson, 2001) was interpreted as support for a potential causal relationship, as it coincides with the inferred divergence time between the western and eastern *D. verneaui* lineages (Macías-Hernández *et al*., 2013). Similar east-west genetic discontinuities coinciding with the limits of the flank collapse within the Anaga peninsula have been found across 13 species of beetle, with genomic evidence for secondary contact and admixture for 12 of the 13 species, and estimated divergence times ranging from ≤0.1 to 4.6 Ma (Salces-Castellano *et al*., 2020). These idiosyncratic divergence times and contemporary admixture, together with more recent evidence for a much older origin for the landslide between 4.2-4.7 Ma (Walter *et al*., 2005) argue against the generality of a causal relationship between mega-landslides and intraspecific divergence. Divergences are more plausibly ascribed to the synergistic role of the landslide-sculpted topography and climatic oscillations throughout the Quaternary forcing distributional shifts across flank collapse limits, promoting cycles of isolation and secondary contact (Salces-Castellano *et al*., 2020, 2021).

It is yet to be understood whether the dynamic of isolation and secondary contact observed by Salces-Castellano *et al*. (2020, 2021) is context-dependent, or if it may apply to other flank collapse systems. More studies are needed that integrate detailed population-level sampling with genetic markers that provide sufficient resolution to allow fundamental predictions from such complex dynamics to be tested. Catastrophic eruptive and erosional activity are likely to be consistent features throughout much of the life cycle of an oceanic island (Jackson, 2013). This in turn suggests a consequential evolutionary impact, and the potential for hypothesis testing when such events are clearly documented in the geological record. One of the best characterised archipelagos, from a geological point of view, is that of the Canary Islands (Carracedo & Troll, 2016). Within this archipelago, the island of Tenerife presents a complex geological history, in which three older volcanic shields are believed to have been originally isolated and then merged within the last 3.5 Ma due to successive volcanic activity (Ancochea *et al*., 1990; Cantagrel *et al*., 1999), or alternatively that two more recent shields formed at the margins of an older and larger central shield, much of which was subsequently overlain with more recent subaerial volcanic activity (Carracedo & Pérez-Torrado, 2013; Carracedo & Troll, 2016; Guillou *et al*., 2004; Figure S1). During the last 2 Ma, Tenerife has suffered several major eruptions (Ancochea *et al*., 1990, 1999; Huertas *et al*., 2002), and has been the subject of many flank collapses (Hunt *et al*., 2014; Figure S1), including some of the largest recorded mega-landslides within the archipelago. The well-documented geological history of Tenerife provides a suitable framework to investigate the impact of geological events on diversification within oceanic islands. Taxa with evidence for recent and ongoing diversification, in turn, constitute fertile ground for investigating the mechanisms promoting divergence among populations.

The *Laparocerus tessellatus* complex of weevil species has been demonstrated to be a suitable model to assess the influence of landscape history on geographic patterns of individual relatedness (García-Olivares *et al*., 2017, 2019). The combination of dispersal limitation within a changing landscape provides a suitable template to investigate how geological dynamics shape intraspecific diversification within an island. A population genomic study within the *L. tessellatus* complex on the island of Gran Canaria has revealed the combined impact of topographic complexity and climate oscillations during the Quaternary on diversification within the complex, in a background of relative geological quiescence (García-Olivares *et al*., 2019). Beyond topoclimatic variation as an engine for diversification, geologically active islands are also likely to structure genetic variation within species (Gübitz *et al*., 2000; Juan *et al*., 2000; Thorpe *et al*., 1996). In contrast to the relative geological dormancy of Gran Canaria over the last 3 Ma, Tenerife represents an island characterised by explosive volcanic activity and numerous gravitational flank collapses over the same period of time. The integration of a fine-scale sampling of the recently diverging *L. tessellatus* complex within the geologically active island of Tenerife thus provides an excellent framework to investigate the role of catastrophic geological activity, in particular mega-landslides, on evolutionary dynamics within islands.

In the present study, we evaluate the evolutionary within-island consequences of mega-landslides using the *L. tessellatus* species complex on Tenerife, which is comprised of two taxonomically described species, *L. tessellatus* Brullé, 1839 and *L. freyi*, Uyttenboogaart, 1940, the latter with four recognised subspecies: *L. f. freyi, L. f. vicarius* Machado 2022, *L. f. punctiger* Machado 2016, and *L. f. canescens* Machado, 2016. Two testable predictions can be made to evaluate the role of mega-landslides on the demographic and evolutionary history within the *L. tessellatus* species complex. The first prediction is that geologically more stable areas within an island may act as reservoirs for population persistence, unlike areas that have suffered catastrophic flank collapses. The second prediction is that areas derived from flank collapses are likely to promote geographical isolation followed by admixture among genomically divergent populations colonising from areas peripheral to the landslide, a process that in turn may result in an increase of genetic diversity under recent or ongoing admixture events (Boca *et al*., 2020). To test these predictions, the *L. tessellatus* complex on Tenerife was sampled representatively across its range, and genome-wide data was integrated into a population genetics and phylogenomics framework to examine the spatial patterns of genetic variation across the island. Specifically, we firstly quantify geographic patterns of genetic diversity and population structure to evaluate their correspondence with contrasting areas of geological stability and flank collapse. Second, we use a simulation-based approach to evaluate competing scenarios of population isolation and estimate timeframes of divergence and gene flow among the inferred genetic groups. These analyses are in turn used to test for temporal congruence with range fragmentation linked to well-documented flank collapses or, alternately, explained by more recent dynamics of population splitting. Finally, we estimate changes in the effective population size through time in order to identify to what extent populations may have responded in parallel to common processes promoting persistence or driving divergence.

## MATERIAL AND METHODS

### Geological context of Tenerife

Tenerife has a complex geological history in comparison with other islands within the Canarian archipelago. During the Miocene, Tenerife is considered to have been composed of three volcanic shields (Figure S1), Roque del Conde (8.9-11.9 Ma), Teno (5.1-6.1 Ma) and Anaga (3.9-4.9 Ma) (Walter *et al*., 2005). These shields are hypothesised to have either been: (i) isolated islands that became fused into the present-day island within the last 3.5 Ma (Ancochea *et al*., 1990; Cantagrel *et al*., 1999), or (ii) to always have been connected via a larger central shield, much of which was subsequently overlain by more recent volcanic material (Carracedo & Pérez-Torrado, 2013). During the last million years, Tenerife has suffered numerous large flank collapses which have left lasting signatures across the landscape (Hunt *et al*., 2014). The prominent scarps along the northern flank of the island represent the mega-landslides of Icod (0.15-0.17 Ma; Masson *et al*., 2002), Orotava (0.54-0.69 Ma; Acosta *et al*., 2003), Roques de García (0.6-1.3 Ma; Acosta *et al*., 2003; Watts & Masson, 1998) and Güímar (0.83-0.85 Ma; Giachetti *et al*., 2011; Hunt *et al*., 2013); the last one in the southern flank (Figure S1; Methods S1).

### Sample collection

Representative geographical sampling from the Tenerife species of the *L. tessellatus* complex was achieved by complementing previous sampling from Faria *et al*. (2016) and García-Olivares *et al*. (2017) with 74 specimens from 61 new localities. This additional sampling effort gave rise to a total of 126 individuals from 102 sites in Tenerife (Table S1). We also included 5 individuals from the *L. tessellatus* complex, belonging to a monophyletic sister clade from the nearby island of Gran Canaria (Garcia-Olivares *et al*., 2019), as an outgroup.

### ddRAD-seq library preparation

We extracted DNA using the Qiagen DNeasy Blood & Tissue kit following the manufacturer’s instructions. DNA was processed using the double-digestion restriction-site associated DNA sequencing protocol (ddRADseq, Peterson *et al*., 2012) as described in Mastretta-Yanes *et al*. (2015) and García-Olivares *et al*. (2019). In brief, DNA was digested with the restriction enzymes MseI and EcoRI (New England Biolabs, Ipswich, MA, USA). Genomic libraries were pooled at equimolar ratios and size selected for fragments between 200-250 base pairs (bp) and, then, sequenced in a single-end 100-bp lane on an Illumina HiSeq2500 platform (Lausanne Genomic Technologies Facility, University of Lausanne, Switzerland).

### Bioinformatic analyses

Raw sequences were demultiplexed, quality filtered and *de novo* assembled using IPYRAD version 0.9.81 (Eaton & Overcast, 2020). Methods S2 provides all details on sequence assembling and data filtering. Unless otherwise indicated (see RAXML and BPP analyses), we performed all downstream analyses using datasets of unlinked SNPs (*i.e*., a single SNP per RAD locus) obtained with IPYRAD considering a clustering threshold of sequence similarity of 0.85 (*clust_threshold*) and discarding loci that were not present in at least 80% individuals (*min_samples_locus*). Optimal parameter values in IPYRAD were identified according to the sensitivity analyses conducted in Garcia-Olivares *et al*. (2019) for the *L. tessellatus* species complex. To exclude the possibility that we had sampled close relatives, we calculated the relatedness between all pairs of genotyped individuals using the *relatedness2* function in VCFTOOLS version 0.1.16 (Danecek *et al*., 2011).

### Genetic clustering analyses

Population genetic structure was assessed using two complementary approaches. First, we used the Bayesian Markov Chain Monte Carlo (MCMC) clustering method implemented in the program STRUCTURE version 2.3.3 (Pritchard *et al*., 2000). We ran STRUCTURE with 200,000 MCMC cycles after a burn-in step of 100,000 iterations, assuming correlated allele frequencies and admixture (Pritchard *et al*., 2000) and performing 40 independent runs for each value of *K* ancestral populations (from *K*=1 to *K*=10). The most likely number of ancestral populations was estimated after retaining the 10 runs per each *K*-value with the highest likelihood estimates. Convergence across runs was assessed by checking the 10 retained replicates per *K*-value provided a similar solution in terms of individual probabilities of assignment to a given ancestral population (*q*-values; Gilbert *et al*., 2012). As recommended by Gilbert *et al*. (2012) and Janes *et al*. (2017), we used two statistics to interpret the number range of ancestral populations (*K*) that best describes our data: log probabilities of Pr(X|*K*) (Pritchard *et al*., 2000) and Δ*K* (Evanno *et al*., 2005), both calculated in STRUCTURE HARVESTER (Earl & vonHoldt, 2012). Finally, we used the Greedy algorithm in CLUMPP version 1.1.2 to align replicated runs of STRUCTURE for the same *K-*value (Jakobsson & Rosenberg, 2007). We also visualised the major axis of genomic variation by performing a Principal Component Analysis (PCA) as implemented in R version 4.0.3 (R Core Team, 2021) using the package *adegenet* (Jombart, 2008). Missing data were replaced by the mean frequency of the corresponding allele estimated across all samples using the ‘*scaleGen’* function (Jombart, 2008).

### Genetic differentiation among individuals

We estimated individual-based genetic distances using Neís distance (Nei, 1972) as implemented in the R package *StAMPP* (Pembleton *et al*., 2013). To assess the relationship among inter-individual genetic distances and geographic distances (isolation-by-distance scenario, IBD), we calculated pairwise weighted topographic distances between each pair of individuals based on a digital elevation model (DEM) at 90-metre resolution using the R package *topoDistance* (Wang, 2020). Methods S3 provides details for the calculation of weighted topographic distances. The matrices were analysed using multiple matrix regressions with randomization (MMRR; Wang, 2013).

### Analysis of genetic diversity and admixture

Heterozygosity is expected to be higher in recently admixed individuals owing to the recombination of source ancestral genomes carrying different genetic variants (Boca *et al*., 2020; Kolbe *et al*., 2008; Witt *et al*., 2023). To test for a positive relationship between heterozygosity and admixture, we calculated observed heterozygosity (*H*_O_) per individual using VCFTOOLS. Individuals were assigned to admixed and non-admixed population groups, according to the probability of assignment (*q*-value) to each ancestral population. Individuals showing a high *q*-value (>95%) to a given ancestral population were assigned as single ancestry, while those showing intermediate *q*-values (5%< *q*-value >95%) were considered to be of admixed origin. Sensitivity analyses were conducted with a more conservative threshold for admixed ancestry (10%< *q*-value >90%), and with admixed individuals divided into two groups to account for the potential effect of varying admixture proportions on genetic diversity (Boca *et al*., 2020). The first group represents more even representation of ancestral population genomes (*q*-values: 30-70%), and the second group more uneven representation (*q*-values: between 5-30% and 70-95%; and alternatively between 10-30% and 70-90%). ANOVAs were used to test for significant differences for *H*_O_ between admixed and non-admixed groups of individuals. To take into account possible geographical clines of genetic diversity (Guo, 2012), relationships between *H*_O_ and spatial variables (longitude and latitude) were tested with linear regressions.

### Phylogenomic inference

Phylogenetic relationships among all individuals were reconstructed using a maximum-likelihood (ML) approach as implemented in RAXML version 8.2.12 (Stamatakis, 2014). Analyses were conducted on a matrix of concatenated SNPs (all SNPs per locus), applying an ascertainment bias correction using the conditional likelihood method (Lewis, 2001). We used a GTR-GAMMA model of nucleotide evolution, performed 100 rapid bootstrap replicates and searched for the best-scoring maximum likelihood tree. Individuals from Gran Canaria were used as an outgroup (Table S1).

We reconstructed phylogenetic relationships among the main genetic groups inferred with clustering analyses, using two coalescent-based methods for species tree estimation: SVDQUARTETS (Chifman & Kubatko, 2014) and SNAPP (Bryant *et al*., 2012). Individuals were assigned to three genetic groups, hereafter referred to as West, North and South, according to the most optimal clustering scheme as inferred in STRUCTURE (*K*=3, see Results section). Only individuals showing the highest probability of assignment (*q*-value >95%) to a given ancestral population Were used for these analyses, resulting in 36 individuals evenly distributed across the three demes. Additional details on SVDQUARTETS and SNAPP analyses are described in Methods S4.

Finally, we used BPP version 4.4.1 (Flouri *et al*., 2018) to estimate the timing of divergence among the three main genetic groups identified by STRUCTURE. Branch length (τ) estimation was performed by fitting the phylogenetic tree inferred in SVDQUARTETS and SNAPP as the fixed topology (option A00 in BPP). These analyses were conducted using the same subset of 36 individuals analysed for group-level phylogenetic inferences. We estimated divergence times using the equation τ=2μ*t*, where τ is the divergence in substitutions per site estimated by BPP, μ is the mutation rate per site per generation, and *t* is the absolute divergence time in years (Walsh, 2001). We assumed a mutation rate of 2.8×10^-9^ calculated for *Drosophila melanogaster* (Keightley *et al*., 2014) which has been estimated to be similar to the spontaneous mutation rate calculated for the butterfly *Heliconius melponeme* (Keightley *et al*., 2015). Details on τ estimation in BPP can be found in Methods S4.

### Inference of past demographic history

We inferred the demographic history of each genetic group using STAIRWAYPLOT2 version 2.1.1, which implements a flexible multi-epoch demographic model on the basis of the site frequency spectrum (SFS) to estimate changes in effective population size (*N*_E_) over time (Liu & Fu, 2020). These analyses were conducted using the subset of 36 individuals with single ancestry assignment. Given a reference genome is not available, we calculated the folded SFS of each genetic group using the script *easySFS.py* (I. Overcast, https://github.com/isaacovercast/easySFS). We considered a single SNP per locus to avoid the effects of linkage disequilibrium. Each genetic group was downsampled to ∼66% of individuals (*i.e*., 8 individuals per deme) to remove all missing data for the calculation of the SFS, minimise errors with allele frequency estimates, and maximise the number of variable SNPs retained. Final SFS for West, North and South contained 13,930, 11,774 and 11,270 variable unlinked SNPs, respectively. Analyses in STAIRWAYPLOT2 were run fitting a mutation rate of 2.8×10^−9^ per site per generation (Keightley *et al*., 2014) and considering a one-year generation time (Machado & Aguiar, 2019). We performed 200 bootstrap replicates to estimate 95% confidence intervals.

### Testing alternative models of divergence and gene flow

We used a simulation-based approach as implemented in FASTSIMCOAL2 version 2.5.2.21 (Excoffier *et al*., 2013) to statistically evaluate the fit of our observed data to alternative scenarios of gene flow between genetic groups and estimate the timing of divergence and gene flow between them (Figure S2). These analyses were conducted using the same subset of 36 individuals analysed for group-level phylogenetic inferences. According to phylogenomic analyses, all scenarios considered an early split between the West genetic group and the ancestor of the North and South genetic groups (T_DIV1_), followed by their divergence (T_DIV2_). Using this topology, we evaluated scenarios assuming (i) divergence in strict isolation, (ii) contemporary gene flow among the three populations and (iii) contemporary gene flow only between West and North, and North and South. We modelled the timing of gene flow as a time interval, represented by a parameter for the time gene flow was initiated (T_MIG1_) and another parameter for the time that it ended (T_MIG2_). In the two isolation-with-migration scenarios, the timing of gene flow (T_MIG1_ and T_MIG2_) was modelled to be fixed across population pairs (models iia and iiia) or, conversely, to vary independently across population pairs (models iib and iiib). We additionally tested the aforementioned isolation-with-migration scenarios without estimating gene flow timing parameters (models iic and iiic). These hypothetical scenarios yielded a total of 7 alternative models (Figure S2). Details on composite likelihood estimation, model selection approach and calculation of confidence intervals for parameter estimates under the most-supported model are described in Methods S5.

## RESULTS

### Genomic data

Illumina sequencing provided a total of 379.54 M sequence reads, with an average of 2.90 M sequence reads per individual (SD=1.74 M) (Figure S3). After the different filtering and assembly steps, each specimen retained on average 50,388 clusters (SD=12,583), with a mean depth per locus of 35.79 (SD=13.58) across individuals. All pairs of genotyped individuals had negative relatedness values (ranging from -3.01 to -0.01), which excludes the possibility that close relatives were included in the analysis (Manichaikul *et al*., 2010).

### Genetic clustering analyses

STRUCTURE analyses identified the most likely number of ancestral populations to be *K*=2 under the Δ*K* criterion (Evanno *et al*., 2005; Figure S4). One of the two ancestral populations was sampled in the westernmost part of the island, the Teno massif, with the other distributed across southern, northern and eastern areas of the island. Individuals with strong signatures of genetic admixture were identified within the northern flank, geographically coinciding with the landslides of Roques García (0.6-1.3 Ma), La Orotava (0.54-0.69 Ma) and Icod (0.15-0.17 Ma) (Figure S5). However, LnPr(X|*K*) steadily increased up to an asymptote of *K*=4. Within this range of likely *K* values, the largest probability gain was from *K*=2 to *K*=3 (Figure S4), suggesting that genetic variation is hierarchically structured within ancestral populations inferred by *K*=2 (Janes *et al*., 2017). Accordingly, when considering *K*=3 genetic variation was further structured into geographically coherent populations hereafter referred to as West, North and South (Figure 1*a*). In a spatial context, individuals inferred to be of single ancestry almost exclusively clustered together geographically, largely within areas that have not been affected by flank collapses, including scarps flanking areas of collapse. Consistent with inferences for *K*=2, individuals of admixed ancestry between West and North were distributed across the aforementioned areas of flank collapse. Additionally within *K*=3, individuals inferred to be of admixed ancestry between East and South were sampled within the geographic limits of the Güímar landslide (0.83-0.85 Ma; Figure 1*a*). These geographic patterns of clustering and admixture persisted when assuming higher *K* values. For *K*=4 and *K*=5, STRUCTURE analyses identified hierarchical population subdivision within the northern population, with individuals assigned with high probability to two ancestral populations, one distributed in the easternmost area of the Anaga peninsula (*K*=4), and the other on the scarps that define the western limits of the Orotava flank collapse (*K*=5; Figure S5). For *K*≥3, individuals of admixed ancestry were consistently found within the La Orotava and Güímar valleys, with a third more limited area of admixture within the western area of the Anaga peninsula with *K*=4 and *K*=5 (Figure S5). Consistent with inferences from STRUCTURE, principal component analysis (PCA) also supported genetic variation to be largely organised into three geographically concordant genetic groups, differentiated across the two first components (Figure 1*b*). Two gradients of admixture were identified within the PCA plot (Figure 1*b*), both corresponding to the two main geographic clines identified in STRUCTURE, each delimited within an area of flank collapse. Differences in allele frequencies between West and all other individuals were largely described along PC1, while differences among individuals from East and South were structured along PC2. Thus, the distributions of individuals from the three ancestral populations within the PCA are congruent with their geographical distributions within the island of Tenerife (Figure 1*a*, *b*).

**Figure 1.**
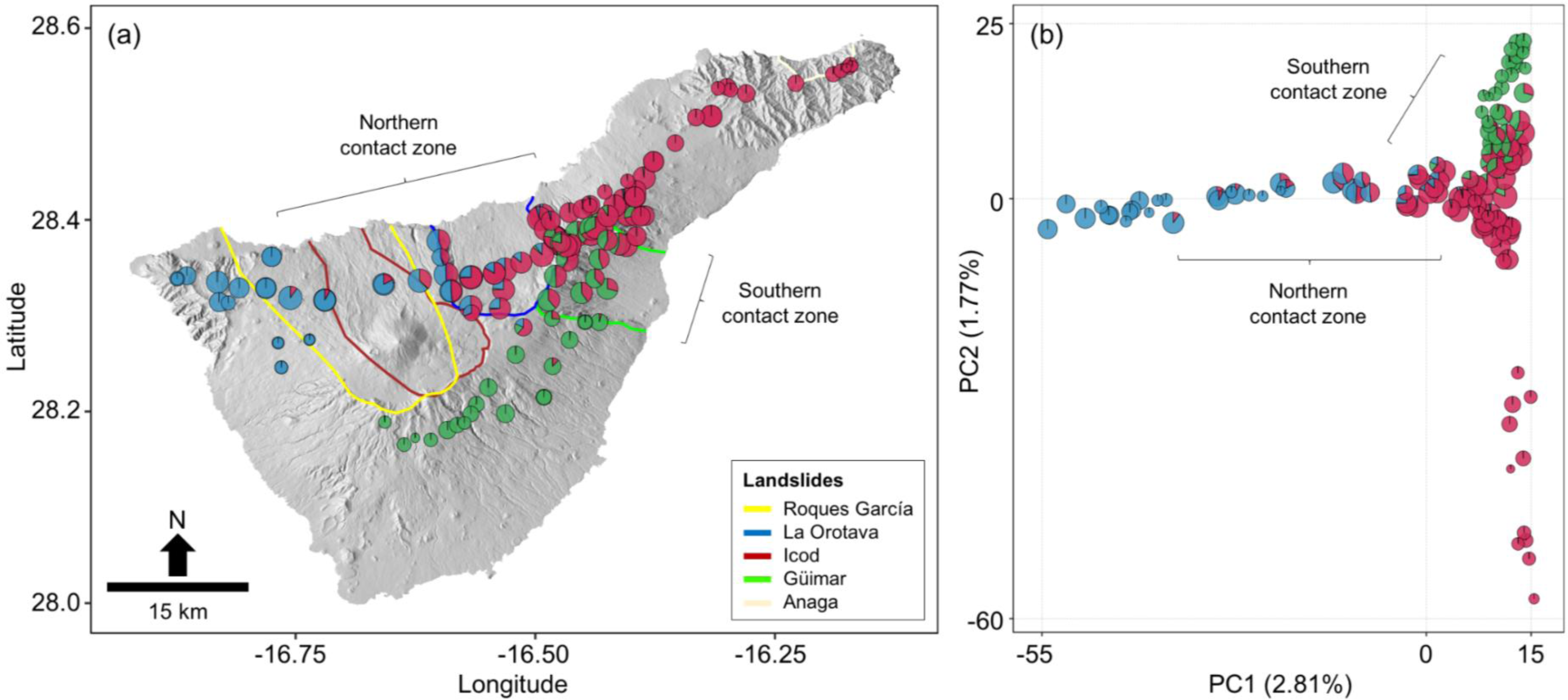
Panel (a) depicts the geographical location of the sampled individuals and their ancestry coefficients (pie charts) as inferred in STRUCTURE, assuming three ancestral populations (*K*=3). Panel (b) shows a principal component analysis (PCA) of genetic variation for the sampled individuals. Pie charts represent the position of each individual along the two first principal components (PCs) and their respective ancestry coefficients based on STRUCTURE results, assuming *K*=3. Size of pie charts in both panels represents individual-based genetic diversity, estimated as observed heterozygosity (*H*_O_).

### Individual-based genetic differentiation, diversity and admixture

Individual-based genetic differentiation estimated with Neís distances were significantly correlated with weighted topographic distances (*R*^2^=0.593, β=0.039, *p*=0.001), consistent with a scenario of isolation-by-distance (IBD). Analyses of average observed heterozygosity (*H*_O_) among individuals of single ancestry (*q*-value >95%) and admixed individuals (5%< *q*-value >95%) revealed significant differences (ANOVA: *F*_4,120_=21.81; *p*<0.001; Figure S6). Individuals derived from both West-North and North-South admixture presented significantly higher *H*_O_ than corresponding single-ancestry populations (*post hoc* Tukey’s tests: *p<*0.001 in all comparisons involving admixed and single-ancestry populations). This result remained significant across all sensitivity analyses (all ANOVAs: *F*_4,120_>11.60; *p*<0.001). Non-parametric Kruskal-Wallis rank sum tests yielded similar results. Finally, *H*o was not significantly associated with latitude or longitude (*p*>0.075), indicating that admixture provides a better explanation for geographic variation in genetic diversity over geographic gradients.

### Phylogenomic inference

The phylogenetic tree reconstructed in RAXML including all individuals revealed the existence of three principal clades corresponding to West, North and South groups, consistent with results from STRUCTURE (Figure S7). The RAXML tree supported a sister relationship between South and North, within a lineage derived from an earlier divergence that gave rise to West. Individuals of admixed ancestry were phylogenetically placed basally within their respective clades (Figure S7).

Analyses in SVDQUARTETS and SNAPP focused on individuals of single ancestry according to STRUCTURE yielded similar inferences. Topologies inferred by both SVDQUARTETS and SNAPP showed fully supported phylogenetic relationships among the STRUCTURE-derived groups and supported an early divergence giving rise to West, followed by a more recent split between North and South (Figure 2), in concordance with the individual-based RAXML analyses. Alternative runs in SNAPP assuming different prior distributions provided similar topologies and branch lengths. According to BPP analyses, the aforementioned splits were estimated to be approximately 0.0033 and 0.0023 τ units (Figure 2). Assuming a mutation rate of 2.8×10^−9^ per site per generation (Keightley *et al*., 2014) and a 1-year generation time (Machado & Aguiar, 2019), both diversification events may have taken place during the Pleistocene, in the Chibanian age, about 0.59 and 0.41 Ma respectively. Inferences of divergence timing from BPP were consistent across runs based on datasets considering different subsets of 5000 loci.

**Figure 2.**
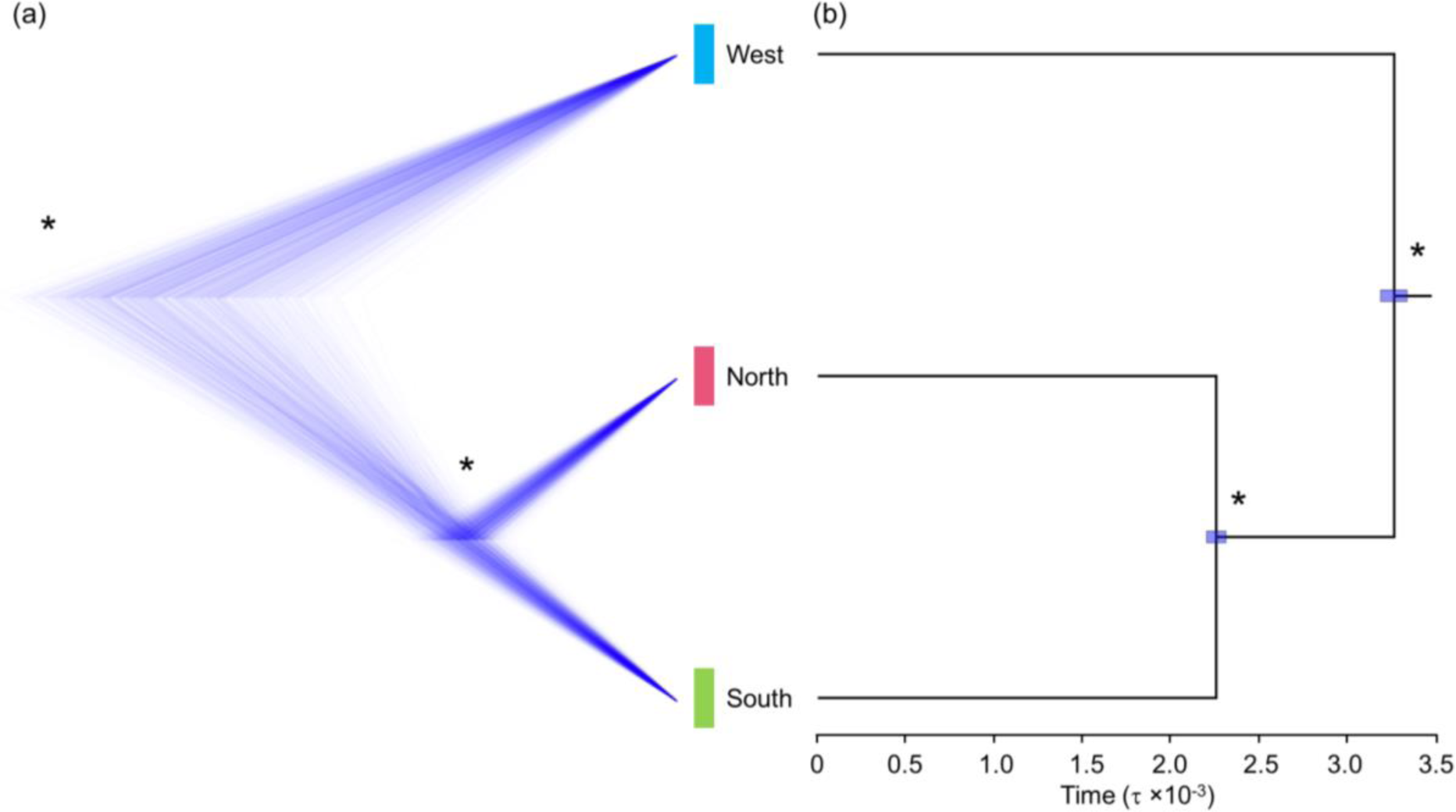
Phylogenetic relationships inferred in SNAPP (panel a) and SVDQUARTETS (panel b) among the three main genetic groups according to STRUCTURE results, assuming three ancestral populations (*K*=3) as the most likely clustering scheme. Panel (b) shows among-group divergence times estimated using BPP with a subset of 5000 randomly chosen loci. The topology was fixed using the phylogenetic tree inferred using SNAPP and SVDQUARTETS. Bars on nodes indicate the 95% highest posterior densities (HPD) of the estimated divergence times. Asterisks denote fully supported nodes in both panels.

### Inference of past demographic history

STAIRWAYPLOT2 analyses indicate that the three genetic groups have experienced parallel demographic responses, undergoing severe demographic declines since the end of the last glacial maximum (LGM, ∼19-21 ka; Figure 3). Consistent with analyses of genetic diversity, West and North presented higher historical estimates of *N*_E_ than South (Figure 3).

**Figure 3.**
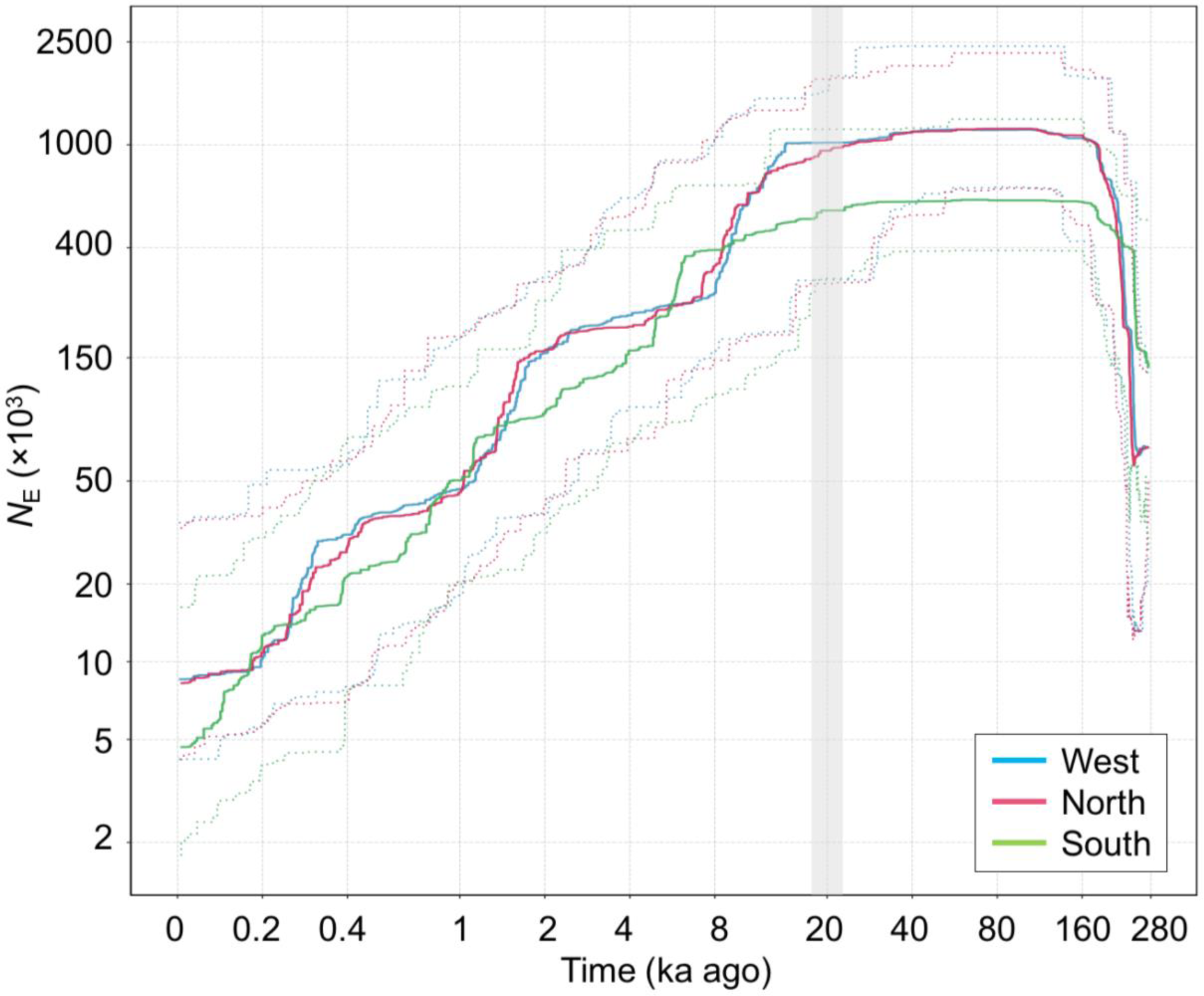
Demographic history of the three main genetic groups as estimated in STAIRWAYPLOT2. Individuals were grouped into the different genetic groups according to the STRUCTURE results assuming three ancestral populations (*K*=3) as the most likely clustering scheme. Continuous and dashed lines represent the median estimate and 95% confidence intervals of the effective population size (*N*_E_), respectively, as inferred in STAIRWAYPLOT2. Axes are logarithmically scaled, with the X axis representing thousands of years (ka). Highlighted period on the X-axis shows the extent of the last glacial maximum (LGM: ∼19-21 ka).

### Testing alternative models of divergence and gene flow

FASTSIMCOAL2 analyses identified the most supported model as being that considering simultaneous gene flow between all population pairs (model iia, Table 1; Figure S2). Considering a 1-year generation time, FASTSIMCOAL2 estimated that all populations diverged from a common ancestor during the Pleistocene, in the Calabrian age, about 1.13 Ma (95% CI: 1.02-1.26 Ma; Figure 4). The posterior divergence between North and South was inferred to take place in the Chibanian age, about 0.50 Ma (95% CI: 0.45-0.56 Ma), consistent with estimates using BPP. Historical gene flow was estimated to be higher both between West and North, and between North and South, compared to between West and South (Figure 4). This coincides with contemporary patterns, with admixture between West and North and between North and South individuals in the Orotava and Güímar valleys respectively, but no evidence for admixture between West and South (Figure S5). Gene flow was estimated to have initiated approximately 154 ka (95% CI: 98-293 ka), considerably after the most recent diversification event (0.50 Ma), and to have ceased approximately 80 ka (95% CI: 26-120 ka; Figure 4). These estimates for the timing of gene flow largely coincide with the time interval where *N*_E_ is estimated to have been higher, according to STAIRWAYPLOT2 analyses (Figure 3).

**Figure 4.**
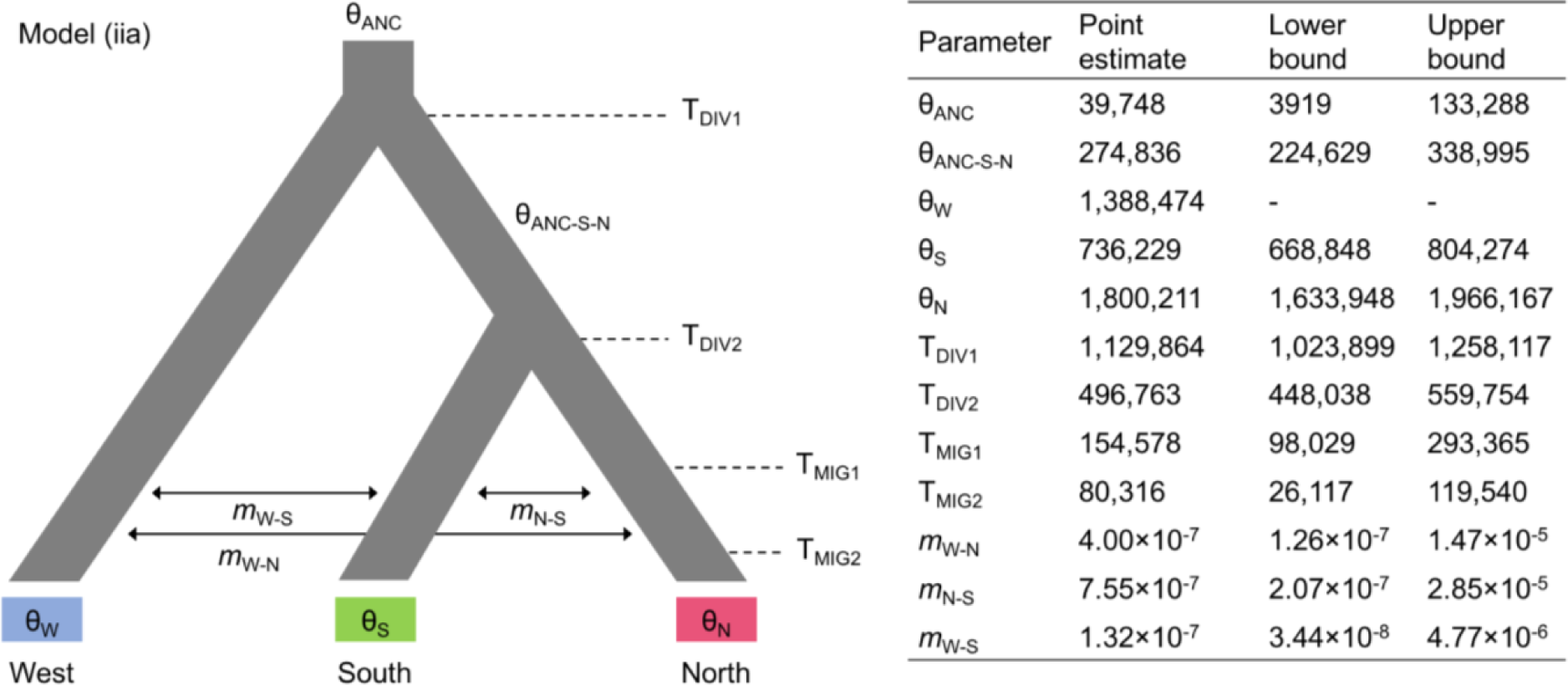
Parameters inferred from coalescent simulations with FASTSIMCOAL2 under the best-supported demographic model. For each parameter, we show its point estimate and lower and upper 95% confidence intervals. Model parameters include ancestral (θ_ANC_, θ_ANC-S-N_) and contemporary (θ_W_, θ_S_, θ_N_) effective population sizes, timing of divergence (T_DIV1_, T_DIV2_), timing of gene flow (T_MIG1_, T_MIG2_) and migration rates per generation (*m*). Note that the effective population size of the West genetic group (θ_W_) was fixed in FASTSIMCOAL2 analyses to enable the estimation of other parameters.

**Table 1.**
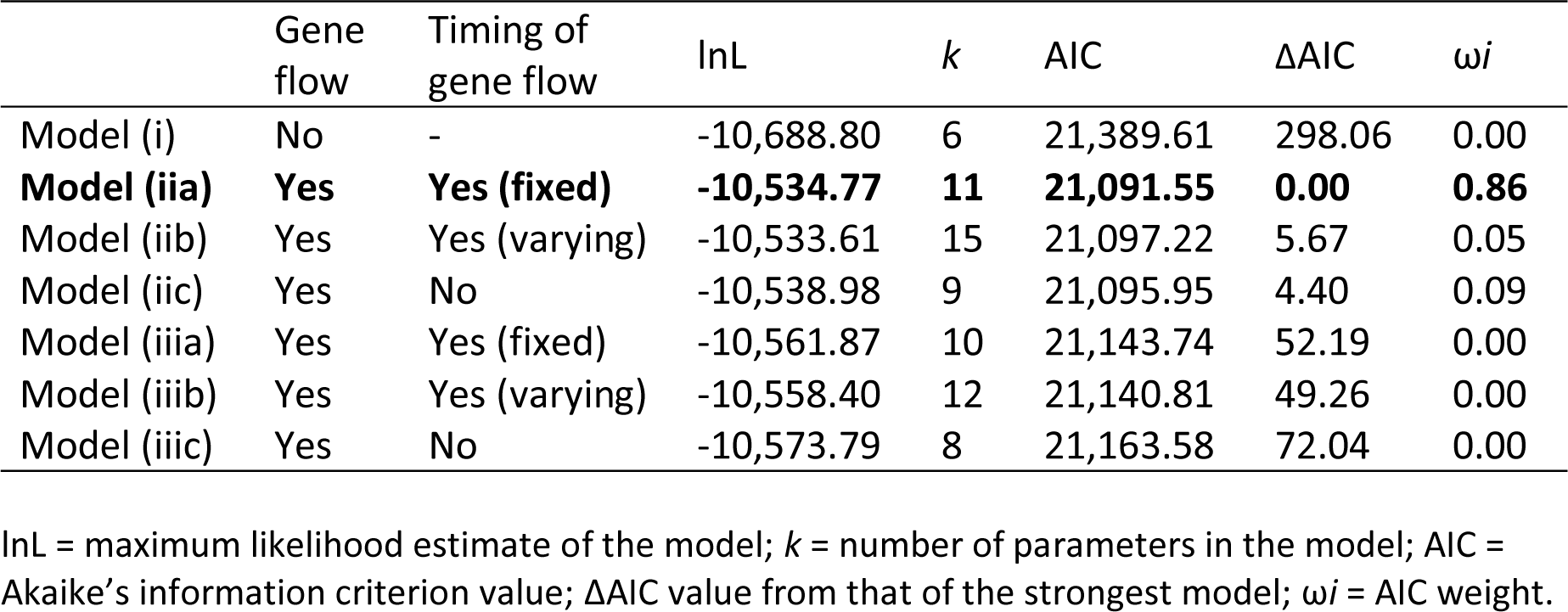
Comparison of alternative models tested using FASTSIMCOAL2 (Figure 4; Figure S2). The best-supported model is highlighted in bold. Models were built both not considering (i) and considering migration among demes (ii and iii). The timing of gene flow was modelled to be fixed across population pairs (iia and iiia) or, conversely, to vary independently across population pairs (iib and iiib). Gene flow was modelled among all demes (ii) or only among West and North, and North and South genetic groups (iii). We also tested scenarios of gene flow without estimating timing parameters (iic and iiic).

## DISCUSSION

It has previously been revealed that mega-landslides can act as drivers for inter-island dispersal of species (García-Olivares *et al*., 2017). However, the evolutionary and demographic consequences of mega-landslides within islands, while having received more interest (Brown *et al*., 2006; Juan *et al*., 2000; Machado, 2022; Macías-Hernández *et al*., 2013), remain less clear. Sampling across a landscape encompassing a sequence of geographically proximate flank collapses, we reveal a dynamic of population isolation and secondary contact that coincides with the landscape features of past flank collapses. In support of our first prediction, we found that individuals with ancestry assignment to a single population were characteristic of relatively geologically stable areas that have neither suffered recent volcanic activity nor flank collapses. The estimated timings of divergence among ancestral populations fall within the geological age estimates for flank collapses within the intervening landscape between areas characterised by ancestral genotypes. In support of our second prediction, individuals with signatures of mixed ancestry were typically sampled within areas of flank collapse. Overall, our study provides a conceptual framework for evaluating the effects of complex geological dynamics in generating novel genetic variation within islands over short spatial scales, through geographic isolation, population persistence and posterior admixture.

### Geological stability, population persistence and differentiation

The geographic distribution of individuals assigned to single ancestral populations within the *L. tessellatus* species complex of Tenerife largely corresponds to areas characterised by long-term geological stability (Figure 1). Hierarchically, three main ancestral populations were consistently inferred in STRUCTURE, of which two were distributed within the northwest and northeast of Tenerife, respectively. These two regions broadly correspond to the Teno massif and the Anaga peninsula (Figure S1), with both regions having remained relatively geologically stable since the end of the Miocene (Carracedo & Pérez-Torrado, 2013), a time interval that encompasses the estimated origin and subsequent diversification of the *L. tessellatus* complex within Tenerife, as inferred in BPP and FASTSIMCOAL2 (see also Faria *et al*., 2016; Machado *et al*., 2017; Machado, 2022). Individuals assigned uniquely to the third ancestral population are almost exclusively associated with areas outside of, but proximate to, scarps defining the Orotava, Roques García and Güímar flank collapse limits. These terrains predate the flank collapses with which they are associated (Figure 1a; Figure S1, Figure S5), thus favouring the persistence of related genomic variation to that extirpated within the areas of flank collapse. Despite uncertainty in divergence time estimates from multiple sources of error (*e.g.*, variation in substitution rate), the timing of divergence among the three ancestral populations (BPP and FASTSIMCOAL2) aligns with the timeframe of the northern flank collapses of Roques de García (0.6-1.3 Ma), Orotava (0.54-0.69 Ma) and Icod (0.15-0.17 Ma), while the split between the South and North populations is estimated to have initiated subsequent to the southern mega-landslide of Güímar (0.83-0.85 Ma; Hunt *et al*., 2014). The geological events across northern Tenerife are suggestive of a cumulative effect on divergence across the geographically proximate and in part overlapping flank collapses of Roques de García, Orotava and Icod.

### Secondary contact and gene flow across areas of flank collapse

While individuals inferred to be of single ancestry were found to be associated with areas of geological stability, a contrasting pattern was observed for individuals of mixed ancestry, which were typically sampled within areas of flank collapse between the ranges of ancestral populations. Estimates of historical gene flow among ancestral populations with FASTSIMCOAL2 indicate that gene flow began approximately 150 ka (95% CI: 98-293 ka) and ceased approximately 80 ka (95% CI: 26-120 ka) (Figure 4). This time interval encompasses the penultimate interglacial period, prior to the onset of the most recent glaciation that culminated 21 ka in the Last Glacial Maximum (LGM), prior to the onset of the current interglacial (Berger *et al*., 2016; Petit *et al*., 1999). More recent and ongoing gene flow is revealed by geographic gradients of admixture between ancestral populations (Verdu & Rosenberg, 2011).

Species within the *L. tessellatus* complex have limited dispersal ability, contributing to geographic structuring of their genetic variation over small spatial scales (García-Olivares *et al*., 2019). While limited dispersal would favour a narrow area of admixture, limited genomic incompatibility among populations and time would favour geographically more extensive gene flow (McEntee *et al*., 2020). Across the combined northern flank collapses of Roques de García, Icod and La Orotava, a gradient of admixture (ancestry assignment to a single population <90%) spans a geographic distance of 25 km. Across the single southern flank collapse of Güímar, a second gradient spans a geographic distance of 10 km. While it remains uncertain when the secondary contact for these two gradients of admixture was initiated, we speculate that it is likely to have been some time after the LGM. Within this temporal window, time since initial secondary contact has been sufficiently long to give rise to geographically extensive admixture within the constraints of limited dispersal. However, higher observed heterozygosity among individuals of admixed origin, compared to individuals of single ancestry, reveals that secondary has been sufficiently recent such that genetic diversity within populations remains above equilibrium expectations (Alcala *et al*., 2013).

### Quaternary climate and species range within flank collapse topography

Analyses with STRUCTURE under increasing values of *K* revealed finer-scale geographic structuring of North with *K*=4 (Figure S5), where genomic variation is organised into western and eastern ancestral populations across the northeastern Anaga peninsula of Tenerife, with geographically intermediate admixed individuals. This pattern was further corroborated by running STRUCTURE with only the eleven individuals sampled in the Anaga Peninsula (Figure S8). This structure coincides with that observed for 13 co-distributed beetle species within the cloud forest of the Anaga peninsula (Salces-Castellano *et al*., 2020), including three related species of *Laparocerus*. Salces-Castellano *et al*. (2021) have revealed that this shared structure across species is best explained by a dynamic of isolation and secondary contact driven by climatic oscillations of the Quaternary. Quaternary climate oscillations within a topographically complex landscape have also been found to explain isolation and secondary contact within the *L. tessellatus* complex on Gran Canaria (García-Olivares *et al*., 2019).

Given the findings of Salces-Castellano *et al*. (2020, 2021) and García-Olivares *et al*. (2019), patterns of admixture across areas of flank collapse within Tenerife are plausibly mediated by range fragmentation and isolation above scarps during glacial climate conditions, with subsequent range expansion and secondary contact during interglacial periods. Further support for such a dynamic comes from demographic reconstructions (Figure 3). In areas occupied by individuals assigned uniquely to one ancestral population, effective population sizes (*N*_E_) are estimated by STAIRWAYPLOT2 to have decreased substantially since the end of the last glacial maximum (LGM: ∼19-21 ka). This generalised response to warming temperatures across all three populations highlights the sensitivity of the focal taxa to climatic variation under a scenario of niche conservatism (Wiens *et al*., 2010).

Globally, climate transition from the LGM until the present is typically characterised by upslope shifts for both the lower and upper elevation limits of species (Davis & Shaw, 2001; Rahbek *et al*., 2019). However, understanding how the distribution limits of the *L. tessellatus* complex may have changed from the LGM until now is complicated by evidence that the lower elevation limits of the orographic cloud bank have been forced downslope since the LGM (Salces-Castellano *et al*., 2021). Given the consistent structuring of genomic variation for the *L. tessellatus* complex in Anaga (Figure S8) with that observed by Salces-Castellano *et al*. (2020), it can reasonably be assumed that lower elevation limits for the *L. tessellatus* complex across the northern slopes of Tenerife have shifted downslope since the LGM. One possibility is that the reduced elevation gradients imposed by scarps when lower elevation limits shift downslope may have facilitated establishment and expansion across the more gradual slopes of flank collapse valley floors. Although the orographic cloud formations are a dominant influence across the northern slopes of Tenerife, elements of laurel forest formations within the scarps of the valley of Güímar (del Arco-Aguilar & Rodríguez-Delgado, 2018) also highlight their potential influence across southern slopes. However, more understanding is needed about local variation in the influence of the orographic cloud layer through time.

## CONCLUSIONS

The paradigm view of the potential evolutionary consequences of mega-landslides on oceanic islands is one of instantaneous range disjunction for species with upper elevational limits that fall below maximum scarp heights, potentially followed by secondary contact. Here we have found support for a model within which the orographic features left by a flank collapse have a more lasting influence on evolutionary processes within species. We reveal that Quaternary climate oscillations can give rise to a cyclical dynamic of range fragmentation and secondary contact across flank collapse landscapes. This dynamic is most likely to be influenced by the sharp elevation gradients associated with scarp height and should be most consequential for species with limited dispersal ability that occupy higher elevations. More generally, our results highlight the role of climate and topography in enhancing genetic diversity within insular species distributions, through both the establishment of divergent populations, and the recombination of their genomes across areas of secondary contact.

## AUTHOR CONTRIBUTIONS

VG-O, JP, HL, VN and BCE conceived the original idea. VN and BCE led the study. VG-O, HL and BCE performed fieldwork. VG-O and YA performed laboratory work with assistance of JP and conducted exploratory analyses. VN conceived the methodological approach and conducted formal analyses. BCE and VN wrote the manuscript. All authors contributed critically to the draft and gave final approval for publication.

## ACKNOWLEDGEMENTS

We wish to thank *Centro de Supercomputación de Galicia* (CESGA) and Teide High-Performance Computing facility (TeideHPC) provided by the *Instituto Tecnológico y de Energías Renovables* (ITER), S.A. for access to computer resources. Fieldwork was supported by the *Cabildo of Gran Canaria* (No. Exp.: 167/15) and *Cabildo of Tenerife* (No. Sigma: 015-00218). This work was supported by the Ministry of Economy and Competitiveness (MINECO) through grants CGL2013-42589-P and CGL2017-85718-P, co-financed by FEDER. VN was supported by a *Juan de la Cierva-Formación* postdoctoral fellowship (FJC2018-035611-I) funded by MCIN/AEI/10.13039/501100011033. VG-O was funded by a FPI pre-doctoral fellowship (BES-2014-067868) from MINECO. JP was supported by the Juan de la Cierva Program-Incorporation (IJCI-2014-19691) and *Ramón y Cajal* Program (RYC-2016-20506) from MINECO.

## DATA ACCESSIBILITY STATEMENT

Upon acceptance, raw Illumina reads will be deposited at the NCBI Sequence Read Archive (SRA) under BioProject xxxxxxx. Input files for all analyses will be available for download from the Dryad Digital Repository (https://doi.org/xx.xxxx/dryad.xxxxxxxxx), upon acceptance. All supplementary tables, figures and methods cited in the main text have been uploaded as Supporting Information.

## BENEFIT-SHARING STATEMENT

All collaborators are included as co-authors in this study, and the results of research have been shared with all relevant parties and the broader scientific community. Benefits from this research accrue from the sharing of our data and results on public databases as described earlier.

## Supporting Information for

**Methods S1.** Geological context of Tenerife

Within the Canary Islands, Tenerife is the largest and highest island, reaching 2034 km^2^ of emerged area and 3,718 meters above sea level. It has a complex geological history in comparison with other islands within the archipelago. The prevailing model for the island of Tenerife argues for the existence of three volcanic shields during the Miocene (Figure S1): Roque del Conde (8.9-11.9 Ma), Teno (5.1-6.1 Ma) and Anaga (3-9-4.9 Ma) (Carracedo & Pérez-Torrado, 2013; Walter *et al*., 2005). These shields are believed to have been originally isolated and then merged within the last 3.5 Ma due to successive volcanic activity (Ancochea *et al*., 1990; Cantagrel *et al*., 1999), or alternatively that the two more recent shields formed at the margins of the older and larger central shield, much of which was subsequently overlain with more recent subaerial volcanic activity (Carracedo & Pérez-Torrado, 2013; Carracedo & Troll, 2016; Guillou *et al*. 2004). As the island reached a higher elevation, the edifice became gravitationally unstable and thus more prone to suffer flank collapses due to volcanic or tectonic seismicity, and dyke injections (McGuire, 2003). As a consequence, Tenerife has suffered numerous large flank collapses, approximately every 150 to 250 ka, with volumes often over 300 km^3^ (Hunt *et al*., 2018). Across the island, 11 documented landslides have left lasting signatures across the landscape (Hunt *et al*., 2014; Watt *et al*., 2014). Vast areas were affected by these mega-landslides, with prominent scarps along the northern flank of the island representing the mega-landslides of Icod (0.15-0.17 Ma; Masson *et al*., 2002), Orotava (0.54-0.69 Ma; Acosta *et al*., 2003), Roques de García (0.6-1.3 Ma; Acosta *et al*., 2003; Watts & Masson, 1998) and Güímar (0.83-0.85 Ma; Giachetti *et al*., 2011; Hunt *et al*., 2013); the last one in the southern flank (Figure S1).

**Methods S2.** Genomic data filtering and sequence assembly

We firstly used FASTQC version 0.11.7 (Andrews, 2010) to quality check raw reads. Then, raw sequences were demultiplexed, quality filtered and de novo assembled using IPYRAD version 0.9.81 (Eaton & Overcast, 2020). Only reads with unambiguous barcodes were retained (*max_barcode_mismatch*) and a stricter filter was applied to remove Illumina adapter contamination (*filter_adapters*). We converted base calls with a Phred score <20 into Ns and discarded reads with >5 Ns (*max_low_qual_bases*). Afterwards, we clustered the retained reads within- and across samples considering a threshold of sequence similarity of 85% (*clust_threshold*) and discarded those clusters with a minimum coverage depth of less than 5 (*mindepth_majrule*). Resulting loci shorter than 35 bp (*filter_min_trim_len*), containing one or more heterozygous sites across more than 50% of individuals (*max_shared_Hs_locus*) and showing more than 20% polymorphic sites (*max_SNPs_locus*) were discarded. In a final filtering step, we only retained loci that were present in at least 80% of the samples (*min_samples_locus*), which yielded a total of 4987 and 6018 unlinked SNPs, when including and excluding the outgroup, respectively. Optimal parameter tuning in IPYRAD is performed following results from the sensitivity analyses conducted by García-Olivares *et al*. (2019) for the *Laparocerus tessellatus* species complex.

**Methods S3.** Weighted topographic distances

We calculated the weighted topographic distances due to Euclidean geographic distances among sampling locations do not likely reflect actual spatial distances among sites in regions of highly topographic complexity (*e.g*., Noguerales *et al*., 2021), such as the island of Tenerife. Topographic distances account for the additional overland distance covered by an organism due to elevation changes imposed by topographic relief. We calculated the weighted topographic distances between each pair of sampled individuals using the ‘*topoWeightedDist’* function from the R package *topoDistance* (Wang, 2020). We calculated the weighted topographic distances using a linear function to weight aspect changes (*hFunction* parameter) and an exponential function to weight the slope (*vFunction* parameter), as recommended by Wang (2020). This assumes that the energetic cost to traverse a slope varies exponentially with the change in angle. We obtained a digital elevation model (DEM) at 30-meter resolution in GeoTIFF format from the EarthExplorer portal (https://earthexplorer.usgs.gov). This DEM is derived from the NASÁs Shuttle Radar Topographic Mission (SRTM) data distributed by the United States Geological Survey (USGS).

**Methods S4.** Phylogenomic inference

Phylogenetic relationships among all individuals were reconstructed using a maximum-likelihood (ML) approach as implemented in RAXML version 8.2.12 (Stamatakis, 2014). Analyses were conducted on a matrix generated by including all SNPs per locus and edited in the R package *phrynomics* (B. Banbury, http://github.com/bbanbury/phrynomics), which resulted in a dataset of 14,928 concatenated SNPs. Accordingly, we applied an ascertainment bias correction using the conditional likelihood method (Lewis, 2001). We used a GTR-GAMMA model of nucleotide evolution, performed 100 rapid bootstrap replicates and searched for the best-scoring maximum likelihood tree. Individuals from Gran Canaria were used as an outgroup (Table S1).

We reconstructed phylogenetic relationships among the main ancestral populations inferred in previous clustering analyses using two coalescent-based methods for species tree estimation: SVDQUARTETS (Chifman & Kubatko, 2014) and SNAPP (Bryant *et al*., 2012). Individuals were assigned to three genetic groups, hereafter referred to as West, North and South, according to STRUCTURE results assuming *K*=3. Only individuals showing the highest probability of assignment (*q*-value >95%) to a given genetic group Were used for this analysis, resulting in 36 individuals evenly distributed across the three genetic groups.

First, we ran SVDQUARTETS (Chifman & Kubatko, 2014) as implemented in PAUP* version 4.0a169 (Swofford, 2002). As for individual-level RAXML analyses, the clade from Gran Canaria was used as an outgroup in SVDQUARTETS (Table S1). We exhaustively evaluated all possible quartets and performed non-parametric bootstrapping with 100 replicates for quantifying topological uncertainty on a dataset of 10,956 unlinked SNPs (including the outgroup).

Second, we ran SNAPP as implemented in BEAST version 2.5.1 (Bouckaert *et al*., 2014). The .*usnps* file from IPYRAD was edited and converted into a SNAPP input file, which resulted in a dataset including 2440 bi-allelic unlinked SNPs shared across branches. Analyses were replicated using different values of the shape (α) and inverse scale (β) parameters of the gamma prior distribution (α=2, β=200; α=2, β=2000) for the population size parameter (θ). The forward (*u*) and reverse (*v*) mutation rates were set to be calculated by SNAPP. We used the log-likelihood correction, sampled the coalescent rate and left default settings for all other parameters. We ran two independent runs for each gamma distribution using different starting seeds for 2 million MCMC generations, sampling every 1000 steps (*ca*. 2000 genealogies). We used TRACER version 1.4 to examine log files, check stationarity and convergence of the chains and confirm that effective sample sizes (ESS) for all parameters were >200. We removed 10% of trees as burn-in and combined tree and log files for replicated runs using LOGCOMBINER version 2.4.7. Maximum credibility trees were obtained using TREEANNOTATOR version 2.4.7 and the full set of retained trees was displayed with DENSITREE version 2.2.6 (Bouckaert, 2010).

Finally, we used BPP version 4.4.1 (Flouri *et al*., 2018) to estimate the timing of divergence among the three main ancestral populations identified by STRUCTURE. The phylogenetic tree inferred in SVDQUARTETS and SNAPP was fitted as the fixed topology in BPP analyses (option A00). These analyses were conducted using the same subset of 36 individuals analysed for group-level phylogenetic inferences. The .*loci* file from IPYRAD was edited and converted into a BPP input file after excluding sequences that were not represented in at least one individual per genetic group using custom R scripts (J-P. Huang, https://github.com/airbugs/; Huang *et al*., 2020). This resulted in a matrix containing 13,714 variable loci. Due to high computational burden, branch length estimation (τ) was inferred using 5000 randomly chosen sequences (for a similar approach see Huang *et al*. 2020). To confirm the consistency of the results, analyses were repeated with four different matrices of 5000 randomly chosen sequences. To ensure convergence, two independent replicates per each locus subset were run for 200,000 generations, sampling every 2 generations, after a burn-in of 40,000 generations. We adjusted the inverse-gamma distributions of θ (α=3, β=0.04) and θ (α=3, β=0.1) priors according to empirical estimates calculated based on the number of segregating sites per site, following Huang *et al*. (2020). We set the “uniform rooted tree” as the species tree prior, applied an automatic adjustment of fine-tune parameters and set the diploid option to indicate that the input sequences are unphased (Flouri *et al*., 2018). We estimated divergence times using the equation τ=2μ*t*, where τ is the divergence in substitutions per site estimated by BPP, μ is the mutation rate per site per generation, and *t* is the absolute divergence time in years (Walsh 2001; *e.g*., Noguerales *et al*., 2021). We assumed a mutation rate of 2.8×10^-9^ calculated for *Drosophila melanogaster* (Keightley *et al*., 2014) which has been estimated to be similar to the spontaneous mutation rate calculated for the butterfly *Heliconius melponeme* (Keightley *et al*., 2015).

**Methods S5.** Testing alternative models of divergence and gene flow

To evaluate the relative statistical support for each of the 7 alternative demographic scenarios (Figure S2), we estimated the composite likelihood of the observed data given a specified model using the site frequency spectrum (SFS) using FASTSIMCOAL2 version 2.5.2.21 (Excoffier *et al*., 2013). We calculated a folded joint SFS using the script *easySFS.py* (I. Overcast, https://github.com/isaacovercast/easySFS). We considered a single SNP per locus to avoid the effects of linkage disequilibrium. Each genetic group was downsampled to ∼66% of individuals (*i.e*., 8 individuals per deme) to remove all missing data for the calculation of the SFS, minimise errors with allele frequency estimates, and maximise the number of variable SNPs retained. The final SFS contained 7,180 variable SNPs. Because we did not include invariable sites in the SFS, we used the “removeZeroSFS” option in FASTSIMCOAL2 and fixed the effective population size (*N*_E_) for West to enable the estimation of other parameters (Excoffier *et al*., 2013; Noguerales & Ortego, 2022; Papadopoulou & Knowles, 2015). We calculated *N*_E_ using nucleotide diversity (π) and estimates of mutation rate per site per generation (μ), where *N*_E_=π/4μ. We estimated π from polymorphic and non-polymorphic loci contained in the .*allele* file from IPYRAD using DNASP version 6.12.03 (Rozas *et al*., 2017). As for previous analyses, we considered a mutation rate per site per generation of 2.8×10^−9^ (Keightley *et al*., 2014).

Each model was run 100 replicated times considering 100,000-250,000 simulations for the calculation of the composite likelihood, 10-40 expectation-conditional maximisation (ECM) cycles, and a stopping criterion of 0.001 (Excoffier *et al*., 2013). We used an information-theoretic model selection approach based on Akaike’s information criterion (AIC) to determine the probability of each model given the observed data (Burnham & Anderson, 2002; *e.g*., Thomé & Carstens, 2016). After the composite likelihood was estimated for each model in every replicate, we calculated the AIC scores as detailed in Thomé and Carstens (2016). AIC values for each model were rescaled (ΔAIC) calculating the difference between the AIC value of each model and the minimum AIC obtained among all competing models (*i.e*., the best model has ΔAIC=0). Point estimates of the different demographic parameters for the best-supported model were selected from the run with the highest maximum composite likelihood. Finally, we calculated confidence intervals (based on the percentile method; *e.g*., de Manuel *et al*. 2016) of parameter estimates from 100 parametric bootstrap replicates by simulating SFS from the maximum composite likelihood estimates and re-estimating parameters each time (Excoffier *et al*., 2013).

---

**Table S1.** Geographic coordinates and elevation (metres above sea level) for each of the individuals from to the *Laparocerus tessellatus* species complex used in this study. The source of each specimen is detailed.

| Island | Population code | Individual code | Elevation | Longitude | Latitude | Source (see footnote on table) |
| --- | --- | --- | --- | --- | --- | --- |
| Gran Canaria | C002 (outgroup) | C002_L123aft | 878 | 28.07254322 | -15.55794230 | Faria *et al.* (2016), García-Olivares *et al.* (2017) |
| Gran Canaria | C003 (outgroup) | C003_L087tir | 870 | 28.06506291 | -15.56382636 | García-Olivares *et al.* (2017) |
| Gran Canaria | C003 (outgroup) | C003_L088tir | 870 | 28.06506291 | -15.56382636 | García-Olivares *et al.* (2017) |
| Gran Canaria | C012 (outgroup) | C012_L309tir | 1556 | 27.98854334 | -15.59328465 | García-Olivares *et al.* (2017) |
| Gran Canaria | C012 (outgroup) | C012_L310tir | 1556 | 27.98854334 | -15.59328465 | García-Olivares *et al.* (2017) |
| Tenerife | T001 | T001_L798spp | 808 | 28.56202187 | -16.17139721 | Faria *et al.* (2016), García-Olivares *et al.* (2017) |
| Tenerife | T002 | T002_L799spp | 768 | 28.56019961 | -16.16919957 | This study |
| Tenerife | T003 | T003_L797spp | 882 | 28.55929840 | -16.17322759 | This study |
| Tenerife | T004 | T004_L159tes | 886 | 28.55855936 | -16.17519236 | Faria *et al.* (2016), García-Olivares *et al.* (2017) |
| Tenerife | T005 | T005_L158tes | 808 | 28.55558312 | -16.18118026 | Faria *et al.* (2016), García-Olivares *et al.* (2017) |
| Tenerife | T006 | T006_L796spp | 788 | 28.55195683 | -16.18922537 | This study |
| Tenerife | T007 | T007_L795spp | 841 | 28.54243006 | -16.22829547 | This study |
| Tenerife | T008 | T008_L161tes | 949 | 28.53192464 | -16.28007027 | Faria *et al.* (2016), García-Olivares *et al.* (2017) |
| Tenerife | T009 | T009_L794spp | 899 | 28.53549911 | -16.29620003 | This study |
| Tenerife | T010 | T010_L146tes | 775 | 28.53869257 | -16.30013311 | Faria *et al.* (2016), García-Olivares *et al.* (2017) |
| Tenerife | T011 | T011_L793spp | 777 | 28.53731314 | -16.30937578 | This study |
| Tenerife | T012 | T012_L613spp | 734 | 28.50833625 | -16.31649694 | This study |
| Tenerife | T012 | T012_L616spp | 734 | 28.50833625 | -16.31649694 | This study |
| Tenerife | T013 | T013_L533spp | 704 | 28.50689194 | -16.33226478 | This study |
| Tenerife | T015 | T015_L469spp | 645 | 28.48002589 | -16.35396053 | This study |
| Tenerife | T016 | T016_L667spp | 910 | 28.46070932 | -16.37668281 | This study |
| Tenerife | T016 | T016_L668spp | 910 | 28.46070932 | -16.37668281 | This study |
| Tenerife | T017 | T017_L607spp | 1121 | 28.44378716 | -16.38643987 | This study |
| Tenerife | T018 | T018_L523spp | 1077 | 28.44044200 | -16.40373413 | This study |
| Tenerife | T019 | T019_L652spp | 1304 | 28.42999464 | -16.39545905 | This study |
| Tenerife | T021 | T021_L375pun | 1290 | 28.42380574 | -16.39610272 | Faria *et al.* (2016), García-Olivares *et al.* (2017) |
| Tenerife | T021 | T021_L376pun | 1290 | 28.42380574 | -16.39610272 | Faria *et al.* (2016), García-Olivares *et al.* (2017) |
| Tenerife | T022 | T022_L090tes | 1464 | 28.42011440 | -16.40749835 | Faria *et al.* (2016), García-Olivares *et al.* (2017) |
| Tenerife | T023 | T023_L528spp | 1106 | 28.42913961 | -16.42743464 | This study |
| Tenerife | T024 | T024_L602spp | 1466 | 28.41454517 | -16.41713273 | This study |
| Tenerife | T025 | T025_L424spp | 1392 | 28.40814756 | -16.40306310 | This study |
| Tenerife | T026 | T026_L428spp | 1300 | 28.40457947 | -16.39634050 | This study |
| Tenerife | T027 | T027_L460spp | 1151 | 28.40373255 | -16.39002131 | This study |
| Tenerife | T028 | T028_L441spp | 1029 | 28.40426000 | -16.38607000 | This study |
| Tenerife | T029 | T029_L430spp | 527 | 28.38284866 | -16.39437182 | This study |
| Tenerife | T030 | T030_L712spp | 1623 | 28.40399285 | -16.42422917 | This study |
| Tenerife | T031 | T031_L672spp | 1737 | 28.39479959 | -16.43164870 | This study |
| Tenerife | T032 | T032_L389spp | 687 | 28.37345156 | -16.40508590 | This study |
| Tenerife | T033 | T033_L449spp | 862 | 28.37479008 | -16.41267523 | This study |
| Tenerife | T034 | T034_L464spp | 1166 | 28.37878915 | -16.42472906 | This study |
| Tenerife | T035 | T035_L404spp | 1728 | 28.39203191 | -16.43735749 | This study |
| Tenerife | T036 | T036_L555spp | 1071 | 28.41501436 | -16.44292499 | This study |
| Tenerife | T036 | T036_L556spp | 1071 | 28.41501436 | -16.44292499 | This study |
| Tenerife | T037 | T037_L642spp | 1700 | 28.39116771 | -16.44141800 | This study |
| Tenerife | T038 | T038_L562spp | 1007 | 28.41115061 | -16.45066119 | This study |
| Tenerife | T039 | T039_L446spp | 1626 | 28.38607142 | -16.44199200 | This study |
| Tenerife | T040 | T040_L662spp | 1730 | 28.38549983 | -16.45585335 | This study |
| Tenerife | T042 | T042_L567spp | 964 | 28.40752049 | -16.46434075 | This study |
| Tenerife | T043 | T043_L169tes | 949 | 28.35900542 | -16.43337187 | Faria *et al.* (2016), García-Olivares *et al.* (2017) |
| Tenerife | T044 | T044_L409spp | 516 | 28.32804785 | -16.42483712 | This study |
| Tenerife | T045 | T045_L403spp | 823 | 28.33886711 | -16.43793667 | This study |
| Tenerife | T046 | T046_L275tes | 1893 | 28.37382685 | -16.46375525 | Faria *et al.* (2016), García-Olivares *et al.* (2017) |
| Tenerife | T047 | T047_L582spp | 1950 | 28.36884949 | -16.46501341 | This study |
| Tenerife | T048 | T048_L692spp | 1771 | 28.37258268 | -16.46783001 | This study |
| Tenerife | T048 | T048_L695spp | 1771 | 28.37258268 | -16.46783001 | This study |
| Tenerife | T049 | T049_L577spp | 1713 | 28.37591345 | -16.47280459 | This study |
| Tenerife | T049 | T049_L578com | 1713 | 28.37591345 | -16.47280459 | This study |
| Tenerife | T049 | T049_L579com | 1713 | 28.37591345 | -16.47280459 | This study |
| Tenerife | T049 | T049_L581com | 1713 | 28.37591345 | -16.47280459 | This study |
| Tenerife | T050 | T050_L147tes | 1468 | 28.38142899 | -16.47936092 | Faria *et al.* (2016), García-Olivares *et al.* (2017) |
| Tenerife | T050 | T050_L148tes | 1468 | 28.38142899 | -16.47936092 | Faria *et al.* (2016), García-Olivares *et al.* (2017) |
| Tenerife | T050 | T050_L273tes | 1468 | 28.38142899 | -16.47936092 | Faria *et al.* (2016), García-Olivares *et al.* (2017) |
| Tenerife | T051 | T051_L268tes | 1174 | 28.39031940 | -16.48924103 | Faria *et al.* (2016), García-Olivares *et al.* (2017) |
| Tenerife | T052 | T052_L597spp | 1044 | 28.39916876 | -16.48384300 | This study |
| Tenerife | T053 | T053_L111tes | 811 | 28.40303852 | -16.49228704 | Faria *et al.* (2016), García-Olivares *et al.* (2017) |
| Tenerife | T054 | T054_L264tes | 752 | 28.40120541 | -16.49567040 | Faria *et al.* (2016), García-Olivares *et al.* (2017) |
| Tenerife | T055 | T055_L279tes | 2010 | 28.35769464 | -16.46666607 | Faria *et al.* (2016), García-Olivares *et al.* (2017) |
| Tenerife | T056 | T056_L261tes | 1050 | 28.36350067 | -16.49300888 | Faria *et al.* (2016), García-Olivares *et al.* (2017) |
| Tenerife | T057 | T057_L283pun | 1941 | 28.34126295 | -16.47890597 | Faria *et al.* (2016), García-Olivares *et al.* (2017) |
| Tenerife | T058 | T058_L414spp | 1142 | 28.32365756 | -16.45175617 | This study |
| Tenerife | T060 | T060_L137pun | 1963 | 28.31639215 | -16.48613927 | Faria *et al.* (2016), García-Olivares *et al.* (2017) |
| Tenerife | T061 | T061_L258tes | 1200 | 28.35539254 | -16.51460599 | Faria *et al.* (2016), García-Olivares *et al.* (2017) |
| Tenerife | T062 | T062_L255tes | 1265 | 28.34795423 | -16.53172068 | Faria *et al.* (2016), García-Olivares *et al.* (2017) |
| Tenerife | T063 | T063_L153tes | 1247 | 28.34422803 | -16.54282731 | Faria *et al.* (2016), García-Olivares *et al.* (2017) |
| Tenerife | T063 | T063_L154tes | 1247 | 28.34422803 | -16.54282731 | Faria *et al.* (2016), García-Olivares *et al.* (2017) |
| Tenerife | T064 | T064_L071fre | 1602 | 28.32716194 | -16.53319716 | Faria *et al.* (2016), García-Olivares *et al.* (2017) |
| Tenerife | T065 | T065_L289fre | 2212 | 28.30758071 | -16.53692393 | Faria *et al.* (2016), García-Olivares *et al.* (2017) |
| Tenerife | T066 | T066_L252fre | 1257 | 28.34006156 | -16.56709586 | Faria *et al.* (2016), García-Olivares *et al.* (2017) |
| Tenerife | T066 | T066_L253fre | 1257 | 28.34006156 | -16.56709586 | Faria *et al.* (2016), García-Olivares *et al.* (2017) |
| Tenerife | T067 | T067_L094fre | 2047 | 28.30291198 | -16.56654087 | Faria *et al.* (2016), García-Olivares *et al.* (2017) |
| Tenerife | T068 | T068_L107fre | 2007 | 28.30911337 | -16.56722321 | Faria *et al.* (2016), García-Olivares *et al.* (2017) |
| Tenerife | T069 | T069_L130fre | 1981 | 28.32568828 | -16.58786492 | Faria *et al.* (2016), García-Olivares *et al.* (2017) |
| Tenerife | T069 | T069_L251fre | 1981 | 28.32568828 | -16.58786492 | Faria *et al.* (2016), García-Olivares *et al.* (2017) |
| Tenerife | T070 | T070_L299fre | 1618 | 28.34281833 | -16.59322012 | Faria *et al.* (2016), García-Olivares *et al.* (2017) |
| Tenerife | T071 | T071_L082fre | 1183 | 28.36095831 | -16.59862291 | Faria *et al.* (2016), García-Olivares *et al.* (2017) |
| Tenerife | T072 | T072_L109tes | 773 | 28.37783460 | -16.60094741 | Faria *et al.* (2016), García-Olivares *et al.* (2017) |
| Tenerife | T073 | T073_L369fre | 1336 | 28.33658376 | -16.62065760 | Faria *et al.* (2016), García-Olivares *et al.* (2017) |
| Tenerife | T074 | T074_L121tes | 1103 | 28.33271243 | -16.65817493 | Faria *et al.* (2016), García-Olivares *et al.* (2017) |
| Tenerife | T074 | T074_L122tes | 1103 | 28.33271243 | -16.65817493 | Faria *et al.* (2016), García-Olivares *et al.* (2017) |
| Tenerife | T075 | T075_L124tes | 1359 | 28.31620437 | -16.71956614 | Faria *et al.* (2016), García-Olivares *et al.* (2017) |
| Tenerife | T075 | T075_L366tes | 1359 | 28.31620437 | -16.71956614 | Faria *et al.* (2016), García-Olivares *et al.* (2017) |
| Tenerife | T075 | T075_L367tes | 1359 | 28.31620437 | -16.71956614 | Faria *et al.* (2016), García-Olivares *et al.* (2017) |
| Tenerife | T076 | T076_L386tes | 502 | 28.36146446 | -16.77505072 | Faria *et al.* (2016), García-Olivares *et al.* (2017) |
| Tenerife | T076 | T076_L387tes | 502 | 28.36146446 | -16.77505072 | Faria *et al.* (2016), García-Olivares *et al.* (2017) |
| Tenerife | T077 | T077_L125tes | 1263 | 28.31853485 | -16.75554197 | Faria *et al.* (2016), García-Olivares *et al.* (2017) |
| Tenerife | T078 | T078_L627spp | 1078 | 28.32877198 | -16.78104466 | This study |
| Tenerife | T078 | T078_L629spp | 1078 | 28.32877198 | -16.78104466 | This study |
| Tenerife | T079 | T079_L134tes | 1018 | 28.32895633 | -16.80887057 | Faria *et al.* (2016), García-Olivares *et al.* (2017) |
| Tenerife | T080 | T080_L069tes | 1245 | 28.31338326 | -16.82018100 | Faria *et al.* (2016), García-Olivares *et al.* (2017) |
| Tenerife | T081 | T081_L475spp | 1146 | 28.31412943 | -16.82984053 | This study |
| Tenerife | T086 | T086_L485spp | 814 | 28.34166966 | -16.86330956 | This study |
| Tenerife | T087 | T087_L489spp | 812 | 28.33847896 | -16.87356573 | This study |
| Tenerife | T087 | T087_L491spp | 812 | 28.33847896 | -16.87356573 | This study |
| Tenerife | T088 | T088_L593spp | 1378 | 28.27159588 | -16.76830880 | This study |
| Tenerife | T088 | T088_L594spp | 1378 | 28.27159588 | -16.76830880 | This study |
| Tenerife | T089 | T089_L557spp | 1207 | 28.24605462 | -16.76495216 | This study |
| Tenerife | T089 | T089_L558spp | 1207 | 28.24605462 | -16.76495216 | This study |
| Tenerife | T090 | T090_L494spp | 1752 | 28.27494931 | -16.73552945 | This study |
| Tenerife | T090 | T090_L496spp | 1752 | 28.27494931 | -16.73552945 | This study |
| Tenerife | T096 | T096_L128can | 2032 | 28.18892526 | -16.65694871 | Faria *et al.* (2016), García-Olivares *et al.* (2017) |
| Tenerife | T097 | T097_L702spp | 1495 | 28.16540488 | -16.63679894 | This study |
| Tenerife | T099 | T099_L509spp | 1676 | 28.17252788 | -16.62524603 | This study |
| Tenerife | T103 | T103_L647spp | 1708 | 28.17034594 | -16.60901984 | This study |
| Tenerife | T104 | T104_L632spp | 1598 | 28.18068328 | -16.59172692 | This study |
| Tenerife | T106 | T106_L722spp | 1579 | 28.18591068 | -16.58144522 | This study |
| Tenerife | T108 | T108_L518spp | 1537 | 28.18815431 | -16.57393027 | This study |
| Tenerife | T110 | T110_L687spp | 1554 | 28.19746381 | -16.56724131 | This study |
| Tenerife | T111 | T111_L682spp | 1624 | 28.20745819 | -16.56140679 | This study |
| Tenerife | T113 | T113_L056can | 1046 | 28.19770092 | -16.53115156 | Faria *et al.* (2016), García-Olivares *et al.* (2017) |
| Tenerife | T115 | T115_L717spp | 1813 | 28.22499033 | -16.54903678 | This study |
| Tenerife | T116 | T116_L419spp | 912 | 28.21490414 | -16.49101821 | This study |
| Tenerife | T116 | T116_L420spp | 912 | 28.21490414 | -16.49101821 | This study |
| Tenerife | T118 | T118_L697spp | 2032 | 28.25930831 | -16.52052323 | This study |
| Tenerife | T119 | T119_L394spp | 1135 | 28.24720547 | -16.48190719 | This study |
| Tenerife | T120 | T120_L543spp | 2187 | 28.28796245 | -16.51246769 | This study |
| Tenerife | T121 | T121_L454spp | 1193 | 28.27461730 | -16.46413989 | This study |
| Tenerife | T122 | T122_L161can | 2016 | 28.29691850 | -16.48256483 | Faria *et al.* (2016), García-Olivares *et al.* (2017) |
| Tenerife | T123 | T123_L163pun | 1383 | 28.29323734 | -16.44797826 | Faria *et al.* (2016), García-Olivares *et al.* (2017) |
| Tenerife | T123 | T123_L164pun | 1383 | 28.29323734 | -16.44797826 | Faria *et al.* (2016), García-Olivares *et al.* (2017) |
| Tenerife | T124 | T124_L165pun | 1078 | 28.29324796 | -16.43328516 | Faria *et al.* (2016), García-Olivares *et al.* (2017) |
| Tenerife | T126 | T126_L800spp | 864 | 28.33485584 | -16.83156566 | This study |

Faria, C.M.A., Machado, A., Amorim, I.R., Gage, M.J.G., Borges, P.A.V. & Emerson, B.C. (2016) Evidence for multiple founding lineages and genetic admixture in the evolution of species within an oceanic island weevil (Coleoptera, Curculionidae) super-radiation. *Journal of Biogeography*, 43, 178-191.
García-Olivares, V., López, H., Patiño, J., Alvarez, N., Machado, A., Carracedo, J.C., Soler, V. & Emerson, B.C. (2017) Evidence for mega-landslides as drivers of island colonization. *Journal of Biogeography*, 44, 1053-1064.

**Figure S1.**
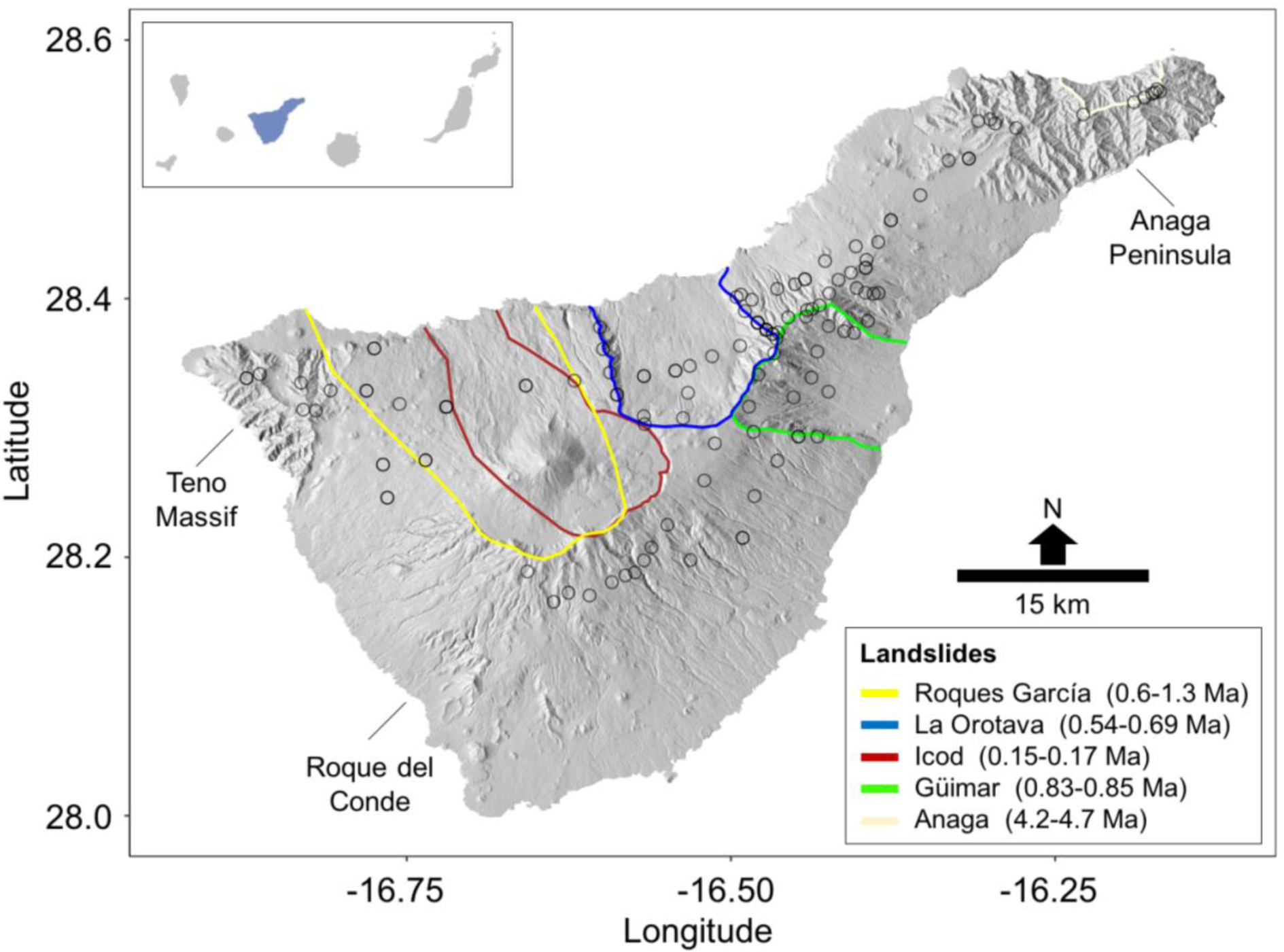
Mega-landslides on the island of Tenerife, which are sufficiently unaffected by subsequent volcanic activity to be able to identify their geographic limits. Open black circles represent sampling sites.

**Figure S2.**
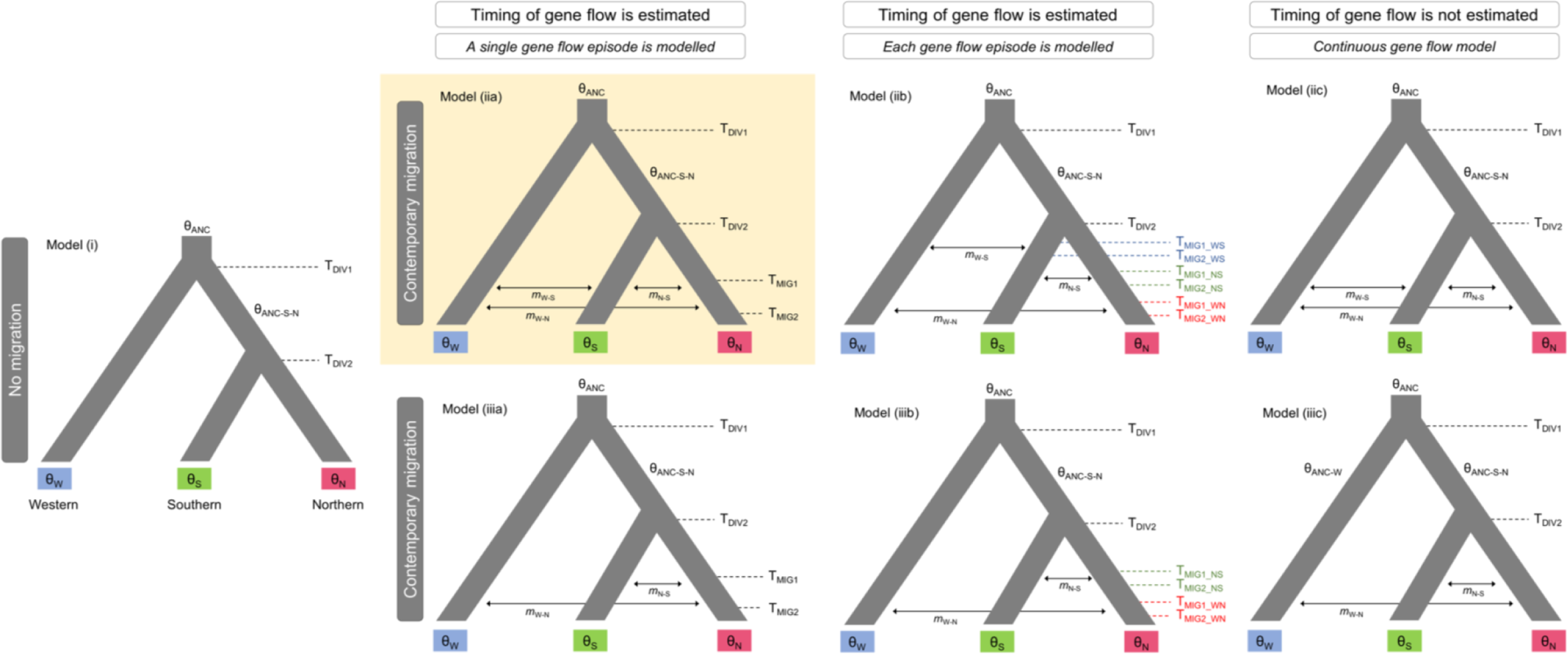
Alternative demographic scenarios tested using FASTSIMCOAL2. Models were tested both considering and not considering contemporary migration. The timing of gene flow was modelled to be fixed across population pairs or, conversely, to vary independently across population pairs. Model parameters include ancestral (θ_ANC_, θ_ANC-W_, θ_ANC-S-N_) and contemporary (θ_W_, θ_S_, θ_N_) effective population sizes, timing of divergence (T_DIV1_, T_DIV2_), timing of gene flow (T_MIG1_, T_MIG2_) and migration rates per generation (*m*). The best-supported model is highlighted.

**Figure S3.**
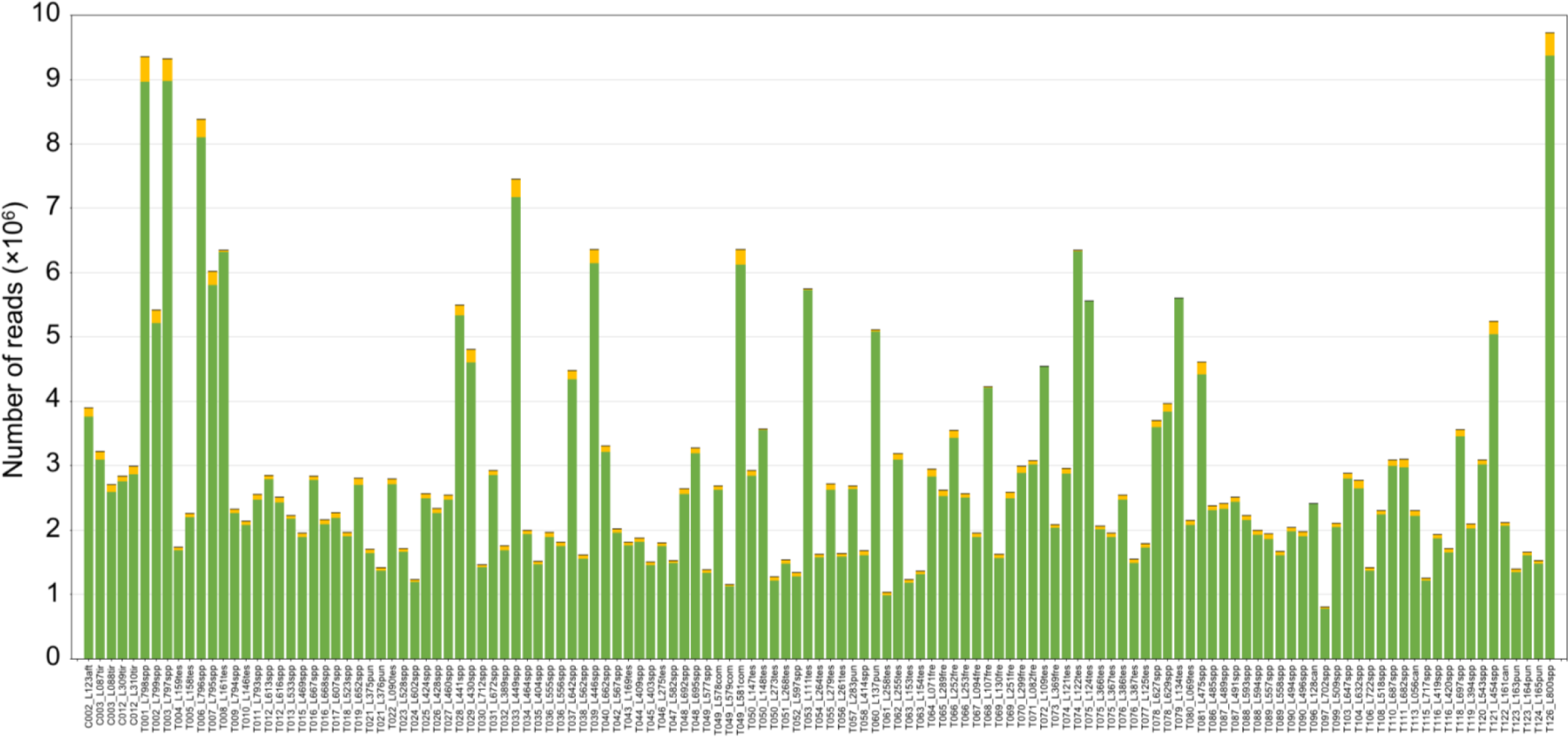
Number of reads per individual before and after different quality filtering steps by IPYRAD. The cumulative stacked bars represent the total number of raw reads obtained for each individual. Within each bar, the orange colour represents the reads that were discarded due to short length (*filter_min_trim_len*). Black colour represents a very small proportion of the reads that were subsequently discarded due to not complying with the quality criteria (*max_low_qual_bases*). Finally, the green colour represents the total number of retained reads used to identify homologous loci during the subsequent steps performed in IPYRAD. Individual codes as in Table S1.

**Figure S4.**
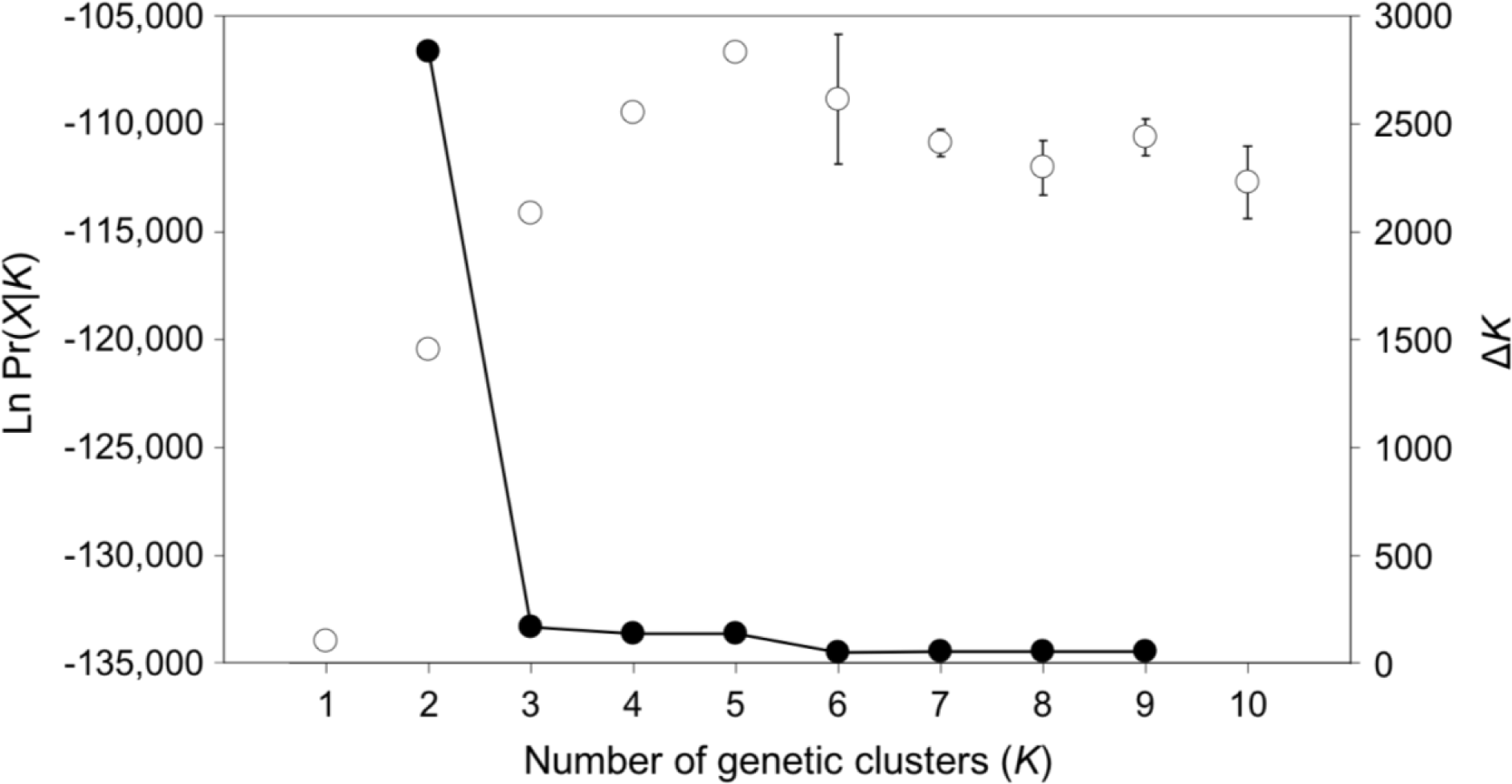
Mean (±SD) log probability of the data (LnPr(X|*K*)) over 10 runs of STRUCTURE (left axes, open dots and error bars) for each value of *K* and the magnitude of Δ*K* (right axes, black dots and continuous line) for analyses based on all ingroup individuals (*n*=126) (Figure 1, Figure S5).

**Figure S5.**
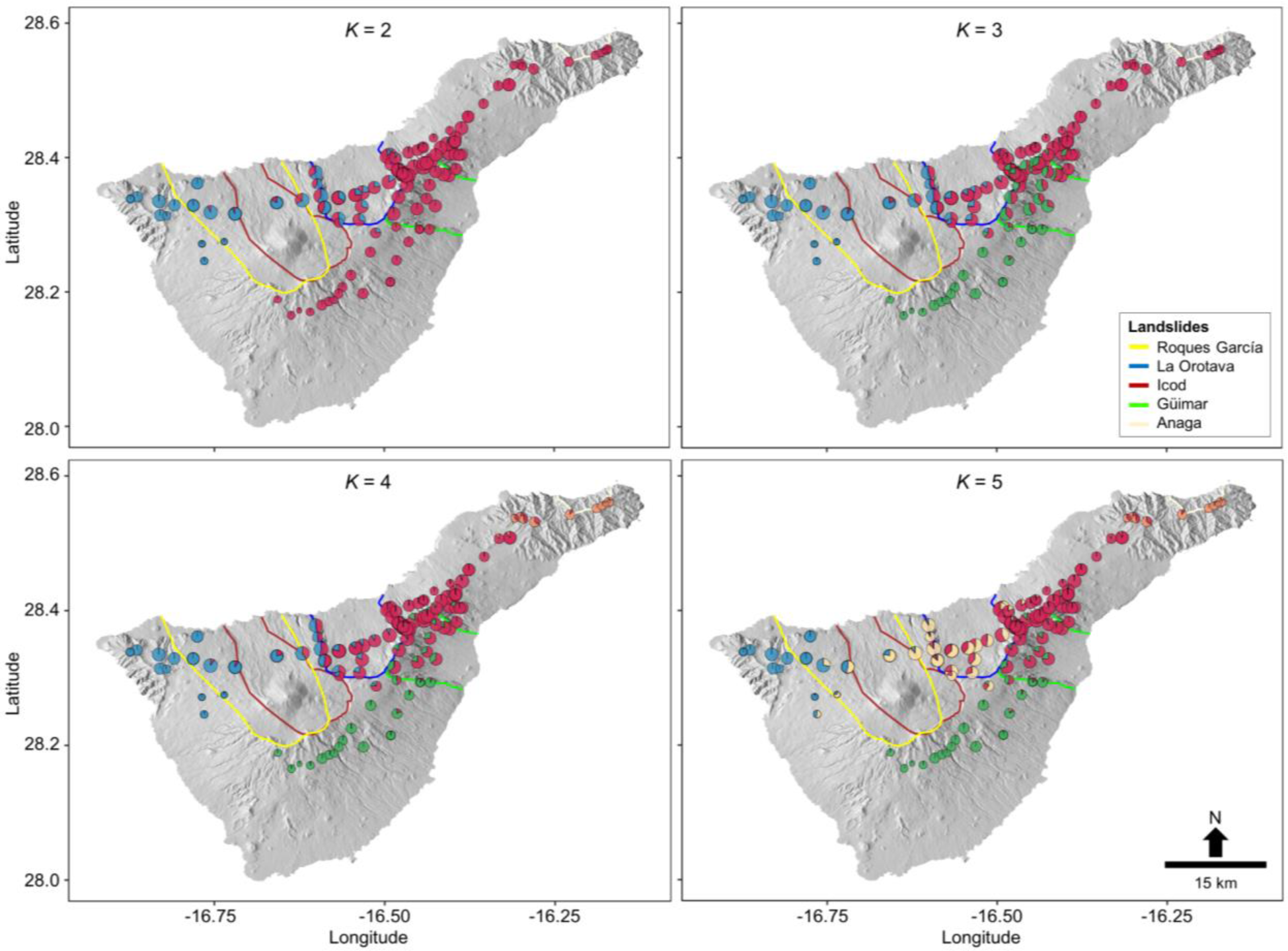
Geographical location of the sampled individuals and their ancestry coefficients (pie charts) as inferred in STRUCTURE, assuming an increasing number of ancestral populations (*K*=2-5). Pie chart size represents individual-based genetic diversity (observed heterozygosity, *H*_O_).

**Figure S6.**
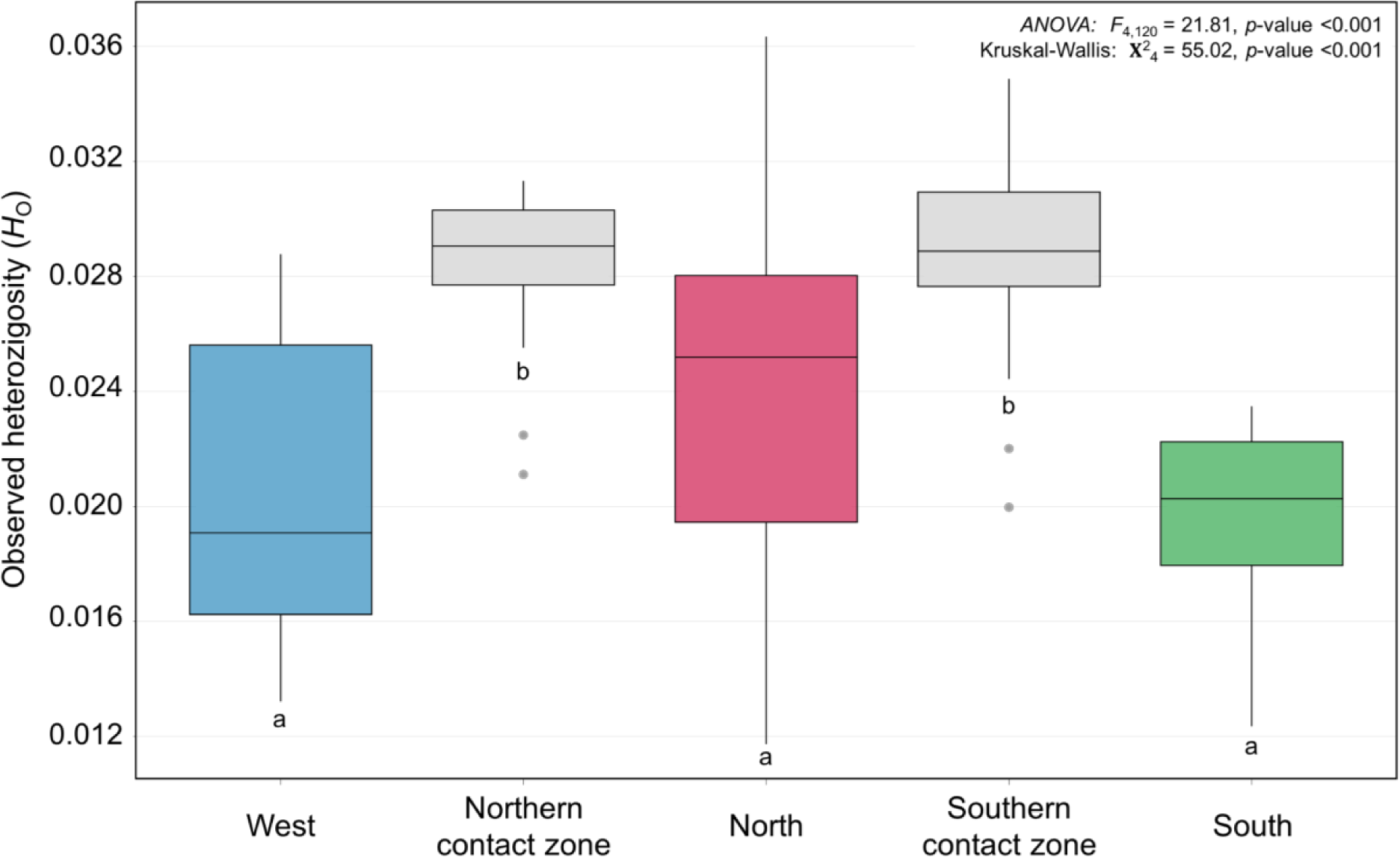
Observed heterozygosity (*H*_O_) across single-ancestry and admixed populations, according to STRUCTURE results, assuming three ancestral populations (*K*=3; Figure 1). Shared letters below the box plots indicate that differences between the respective groups are not statistically significant (*p*-value >0.05) after *post hoc* tests using both the Tukey and Wilcoxon methods.

**Figure S7.**
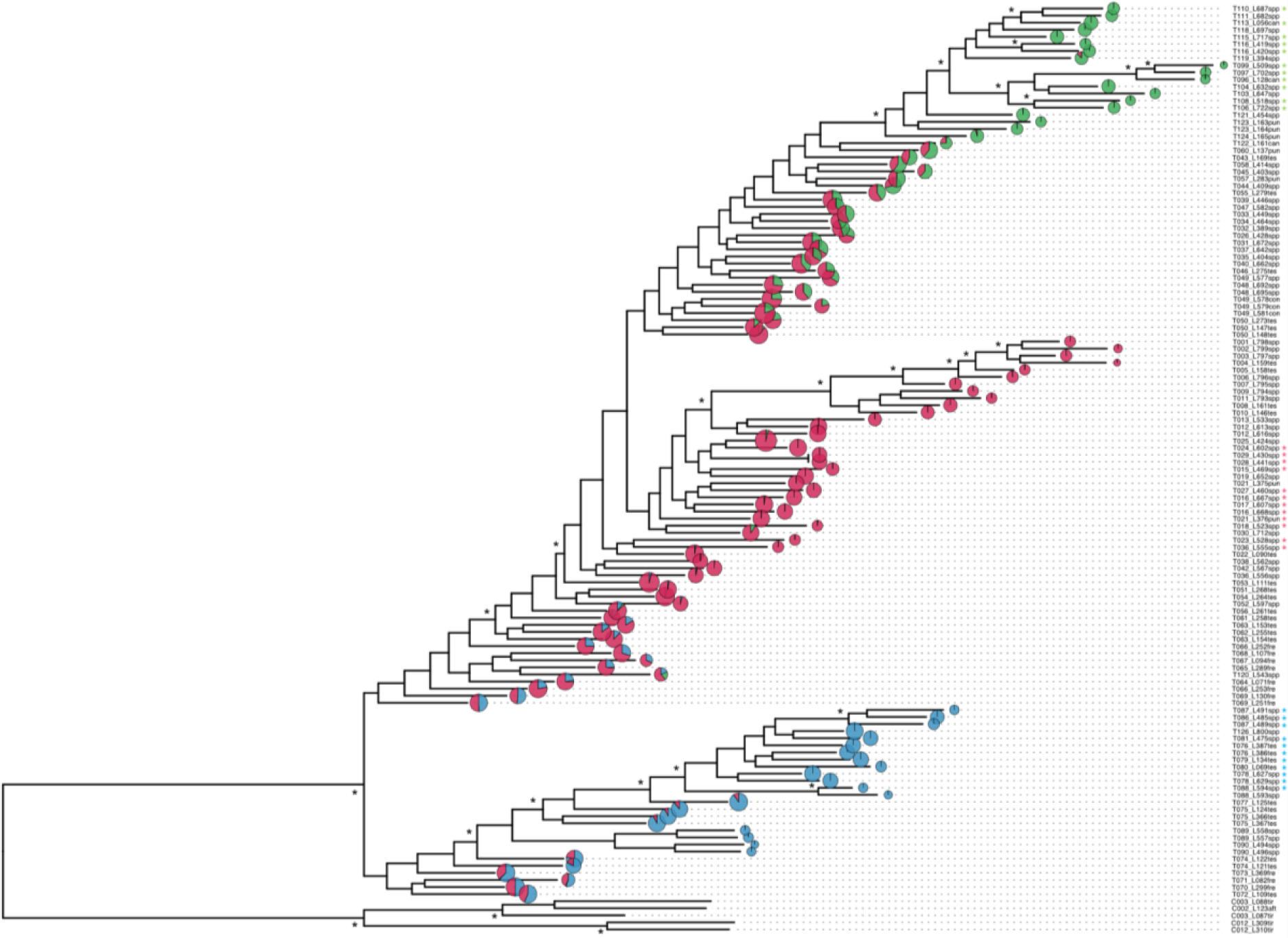
Phylogenetic relationships among individuals as inferred in RAXML. Asterisks on the tree denote well supported-nodes (bootstrap value >90). Pie charts represent the individual-based ancestry coefficients as inferred STRUCTURE, assuming three ancestral populations (*K*=3). Size of pie charts represents individual-based genetic diversity (observed heterozygosity, *H*_O_). Five individuals belonging to a monophyletic sister clade from the nearby island of Gran Canaria were used as an outgroup. Colour asterisks at the right of the individual codes represent those specimens used for analyses in BPP, STAIRWAYPLOT2 and FASTSIMCOAL2. Individual codes as in Table S1.

**Figure S8.**
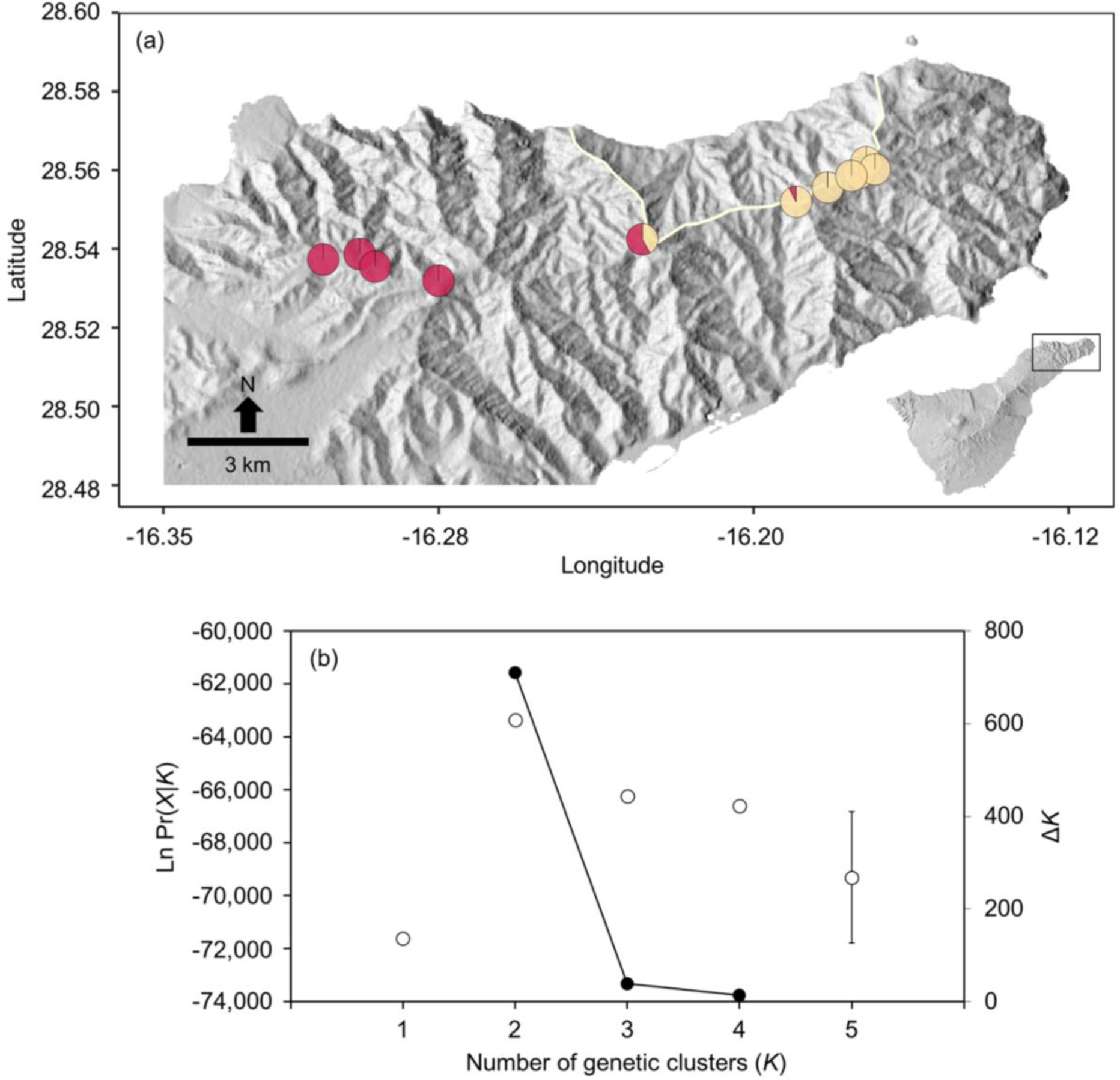
Panel (a) depicts the geographical location of the individuals sampled in the Anaga Peninsula and their ancestry coefficients (pie charts) as inferred in STRUCTURE, assuming two ancestral populations (*K*=2). Geographic boundaries of the Anaga landslide are shown in light yellow. Panel (b) represents the mean (±SD) log probability of the data (LnPr(X|*K*)) over 10 runs of STRUCTURE (left axes, open dots and error bars) for each value of *K* and the magnitude of Δ*K* (right axes, black dots and continuous line) for analyses based exclusively on the eleven individuals sampled in the Anaga Peninsula.

